# Deep attention networks reveal the rules of collective motion in zebrafish

**DOI:** 10.1101/400747

**Authors:** Francisco J.H. Heras, Francisco Romero-Ferrero, Robert C. Hinz, Gonzalo G. de Polavieja

**Affiliations:** Champalimaud Research, Champalimaud Centre for the Unknown, Lisbon, Portugal

## Abstract

A variety of simple models has been proposed to understand the collective motion of animals. These models can be insightful but lack important elements necessary to predict the motion of each individual in the collective. Adding more detail increases predictability but can make models too complex to be insightful. Here we report that deep attention networks can obtain in a data-driven way a model of collective behavior that is simultaneously predictive and insightful thanks to an organization in modules. The model obtains that interactions between two zebrafish, *Danio rerio*, in a large groups of 60-100, can be approximately be described as repulsive, attractive or as alignment, but only when moving slowly. At high velocities, interactions correspond only to alignment or alignment mixed with repulsion at close distances. The model also shows that each zebrafish decides where to move by aggregating information from the group as a weighted average over neighbours. Weights are higher for neighbours that are close, in a collision path or moving faster in frontal and lateral locations. These weights effectively select 5 relevant neighbours on average, but this number is dynamical, changing between a single neighbour to up to 12, often in less than a second. Our results suggest that each animal in a group decides by dynamically selecting information from the group.

**Highlights:**

- At 30 days postfertilization, zebrafish, *Danio rerio*, can move in very cohesive and predictable large groups
- Deep attention networks obtain a predictive and understadable model of collective motion
- When moving slowly, interations between pairs of zebrafish have clear components of repulsion, attraction and alignment
- When moving fast, interactions correspond to alignment and a mixture of alignment and repulsion at close distances
- Zebrafish turn left or right depending on a weighted average of interaction information with other fish, with weights higher for close fish, those in a collision path or those moving fast in front or to the sides
- Aggregation is dynamical, oscillating between 1 and 12 neighbouring fish, with 5 on average

## Introduction

There is a wide range of models of collective behavior. A useful way to understand the relative merits of these models is to classify them by their accuracy and their complexity (e.g. [1, 2]). Some of the classical models of collective behavior, like interaction models [3, 4, 5, 6, 7], many-eyes or weighted averages [8, 9, 10, 11], Condorcet [12] or others [13, 14, 15, 16, 17] are of very low complexity. Low complexity can be formally characterised [18], but in practical terms we can define it as the number of parameters in the model. If a model has a low parameter-complexity, we can write down the mathematical description and study it in detail, leading to an intuitive grasp of the problem and therefore a better design of new experiments. In general, however, this simplicity likely misses important biological components. For this reason, these low-parameter-complexity models are not typically tested in their detailed quantitative predictions, using simpler global parameters instead (but see [19, 20]).

New techniques allow to individually track each animal in large collectives with high precision [21], and can provide enough data to build accurate models of collective behaviour. However, it is difficult to increase accuracy without increasing complexity. For example, we can use our trajectories to train a very precise model base on deep neural networks, because they contain thousands or millions of parameters that can be adjusted to approximate any possible function [22]. Unfortunately, this parameter-complexity typically makes deep neural networks black boxes difficult to analyse. However, a function that is parameter-rich but has few inputs and outputs (i.e. low variable-complexity) can in principle be understood through graphical plotting. Here we propose to use deep attention networks [23, 24, 25], because they express the social interaction as a combination of two deep-network modules of few inputs and outputs each, and thus they allow to simultaneously achieve insight and predictive accuracy.

We can illustrate the reduction of variables in a modular model by comparing it against an equivalent non-modular model. Without a modular approach, a case with, say, 25 neighbours, and with each animal being sensitive to the 2D position, speed and orientation of each, would need to be modelled using 25 × 6 = 150 variables for each individual. Studying a problem in 150 dimensions is impractical as insight is very difficult to extract. However, deep attention networks [23, 24, 25] are organized in modules, and for some problems each module might depend only on a small number of variables, of the order of 4 − 6 in our problem. This reduction from 150 variables to modules of 4 − 6 variables allows for insight into the rules by analysis of these these modules, while achieving high prediction accuracy.

## Results

### Predicting the future using a deep interaction network

We recorded videos of groups of 60, 80 or 100 juvenile zebrafish, *Danio rerio*, (Figure 1A for a detail; see [21] for setup). We tracked videos using our system idtracker.ai, obtaining high-quality position, velocity and acceleration values (see **Methods and Materials**).

**Figure 1:**
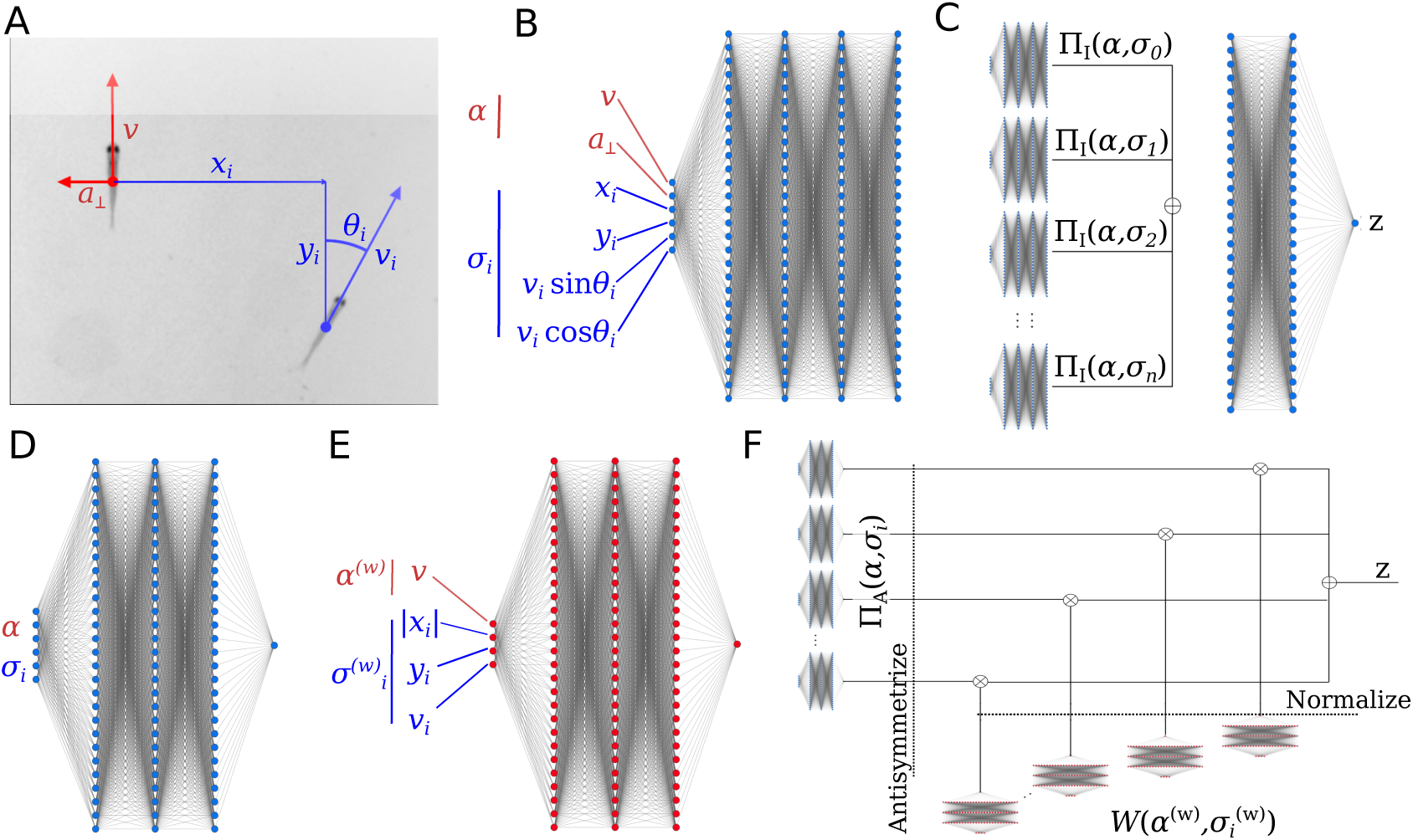
Deep-learning a model of collective behaviour. (**A**) Variables used to predict future turns. Asocial variables, those only involving the focal, in red. Social variables, those involving both the focal and a neighbour, in blue. (**B**) Pair-interaction subnetwork, receiving asocial variables *α* and social variables *σ_i_* from a single neighbour *i*, and outputting a vector of 128 components. All pair-interaction networks share the same weights. (**C**) Interaction network, showing how the outputs of the pair-interaction subnetworks, one for each neighbour, are summed and then fed to an interaction subnetwork. The output, *z* is the logit of the focal fish turning right after 1 s. (**D**) Pair-interaction subnetwork of the attention network. (**E**) Aggregation subnetwork of the attention network. Same structure as D, but the input is a restricted symmetric subset of the variables and the output is passed through an exponential function to make it positive. (**F**) Attention network, showing how the inputs of the pair-interaction and attention subnetworks are integrated to produce a single logit *z* for the focal fish turning right after 1 s.

We used the trajectories to obtain data-driven models of fish interactions. First, we required our models to be predictive of the future of a focal fish in test data (video sequences not used to train the model). The requirement of biological insight, which we discuss in the next section, was only added later. The reason for this strategy is that we first need to find out how much we can predict from video and models designed to be insightful need to make assumptions that may reduce the ability to predict.

We used a deep interaction network, inspired by their success in interacting systems in Physics [26]. Our deep interaction network is divided in two parts: (i) *n* pair-interaction subnetworks, each describing the interaction of a focal fish with one of its *n* closest neighbors (Figure 1B), and (ii) an aggregation or weighting subnetwork, aggregating the *n* outputs of the pair-interaction subnetworks (Figure 1C, subnetwork to the right).

The inputs to the network are quantities expressed in a coordinate system centered at the focal fish and with the *y*-axis in the direction of the velocity of the focal (Figure 1A, red). The pair-interaction subnetwork has as inputs the asocial information of the focal, *α* (Figure 1B, red), and the social information of one neighbour *i, σ_i_* (Figure 1B, blue). The asocial information of the focal is its speed, *v*, tangential acceleration, *a*_||_, and normal acceleration, *a*_⊥_. We found that *a*_||_ had little impact on accuracy (Table S1), so we did not consider it in further computations. The social information is the neighbour position with respect to the focal, *x_i_* and *y_i_*, its velocity, *v_i,x_* and *v_i,y_*, and acceleration, *a_i_,_x_* and *a_i,y_*. Neighbour accelerations had little impact on accuracy (Table S1) and were not used in further computations.

Accuracy of prediction of the turning side of the focal fish after 1 second evaluated on held-out test data improves with the number of neighbours, but with diminishing returns (Figure S1); we chose *n* = 25 neighbors. In the main text we provide analysis of groups of 100 animals and prediction at 1 s in the future for illustration purposes. Our models predict well a range of futures (Figure S2). Results on how fish interact were found to be similar in computations using 250 ms, 500 ms and 1.5 s in the future (Figure S3, Figure S4, Figure S5) and for groups of 60 or 80 zebrafish (Figure S6, Figure S7).

Accuracy of prediction of the turning side after 1 s is higher for large turning angles than for turning angles close to 0 or 180 degrees (Figure S8). For turning angles of 20 – 160°, the interaction network predicted the correct side with an accuracy of 84.4%; up to 87.1 % for 30 – 100°. In contrast, a model using only focal variables failed to obtain a high accuracy and reached only 55 %. Interaction networks with different architectures performed slightly worse (Table S2), while fully-connected networks performed consistently worse (Table S3).

The high accuracy of the interaction network shows that the 6 × 25 = 150 dimensions capture an important part of the collective dynamics. The instances not predicted may originate from a variety of effects, including higher-order correlations, individuality and non-markovian effects, i.e. history-dependency, be it at short scales or at long scales (internal states or unaccounted behavioural variables, like posture or eye movements). Accuracies were larger the larger the group (Table S4), consistent with the idea that interactions lock individuals into social dynamics, less stochastic and less dependent on individual internal states than asocial dynamics.

### Deep attention networks obtain a predictive and analyzable model

The deep interaction model of the previous section taught us which are the relevant variables to consider and gave us a reference accuracy. However, the model is too high-dimensional to provide useful insight into animal interactions. The pair interaction subnetwork, for example, takes the values of 6 variables as inputs and outputs 128 values (Figure 1B). The aggregation subnetwork first sums up the 25 128-dimensional vectors to give a single vector of 128 components, and then processes it to output a single number, *z* (Figure 1C).

To gain insight on the nature of fish interactions, we used a deep attention network [23, 24]. Like the interaction network, the attention network has a subnetwork to describe the interaction of the focal with each of the *n* neighbors, except now with a single output (Figure 1D). The aggregation subnetwork is a function weighting differently each neighbor depending on its kinematic parameters and those of the focal (Figure 1E). We found that focal and neighbor speed and neighbor position are the inputs to the aggregation subnetwork with the highest impact in accuracy. We can express the probability that the focal turns to the right after 1 s, *p*, as *p* = 1/1 + exp(− *z*), where z is the logit that the deep attention network outputs (Figure 1F),

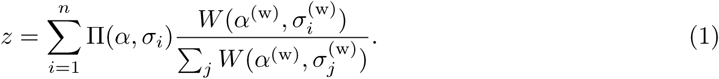

The pair-interaction subnetwork, Π(*α*, *σ_i_*), describes the interaction of the focal and one neighbour *i*. The aggregation subnetwork *W* gives different weights to the different neighbors *i* in the aggregation depending on the kinematic parameters of focal *α*^(^*^w^*^)^ and neighbor relative to focal 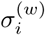. The subscript indicates that these variables may be different to the ones in the pair-interaction subnetwork.

Since we want the pair-interaction subnetwork, Π, and the aggregation subnetwork, *W*, to represent the logit of turning after 1 s given a neighbor and a weight representing the importance of that neighbor, respectively, they must differ on several accounts. (i) Π can have any real output, while *W* must be always positive. (ii) Π must be antisymmetric with respect to reflection on the y-axis, while *W* must be symmetric. This is because we assume that a neighbour to the right makes the focal go to the right as much as an identical neighbour to the left of the focal makes the focal move to the left. For the aggregation weight *W*, however, we assume that the importance of the two cases is the same. (iii) The aggregation weights must sum 1. These three conditions are required and we enforced them by: (i) using an exponential as final activation function of *W*, (ii) antisymmetrizing Π and using symmetric input in *W*, and (iii) normalising the outputs of *W* by the sum across all neighbours prior to the integration with the outputs of Π.

The resulting attention network achieves 83.2% accuracy for turns between 20° and 160° and around 85.9% for 30-100°. This is slightly less accurate than the interaction network, but the much lower dimensionality of the two subnetworks allows for a detailed analysis.

### The structure of interaction of a pair of animals in a collective

The pair-interaction subnetwork Π is a six-dimensional function. We plotted its output, the logit of the focal fish turning to the right after 1 s, *z*, as a function of two variables: the angle, *θ_i_*, and the speed of the neighbour, *v_i_* (Figure 2). We fixed the other four variables: focal at median velocity of 3.04 BL/s, focal normal acceleration *a*_⊥_ = 0, and neighbour position at *x_i_* = 7 BL and *y_i_* = 1 BL. At a neighbour velocity above the median (Figure 2A, left; median speed indicated with a horizontal line at 3.04 BL/S), the focal animal is sensitive to the neighbour orientation, with a high probability of turning right (left) after 1 s when the neighbour is moving away from (toward) the focal, resulting in an alignment of the focal to the neighbour. When the neighbour speed is below the median, however, the focal is attracted towards it regardless of the neighbour orientation.

**Figure 2:**
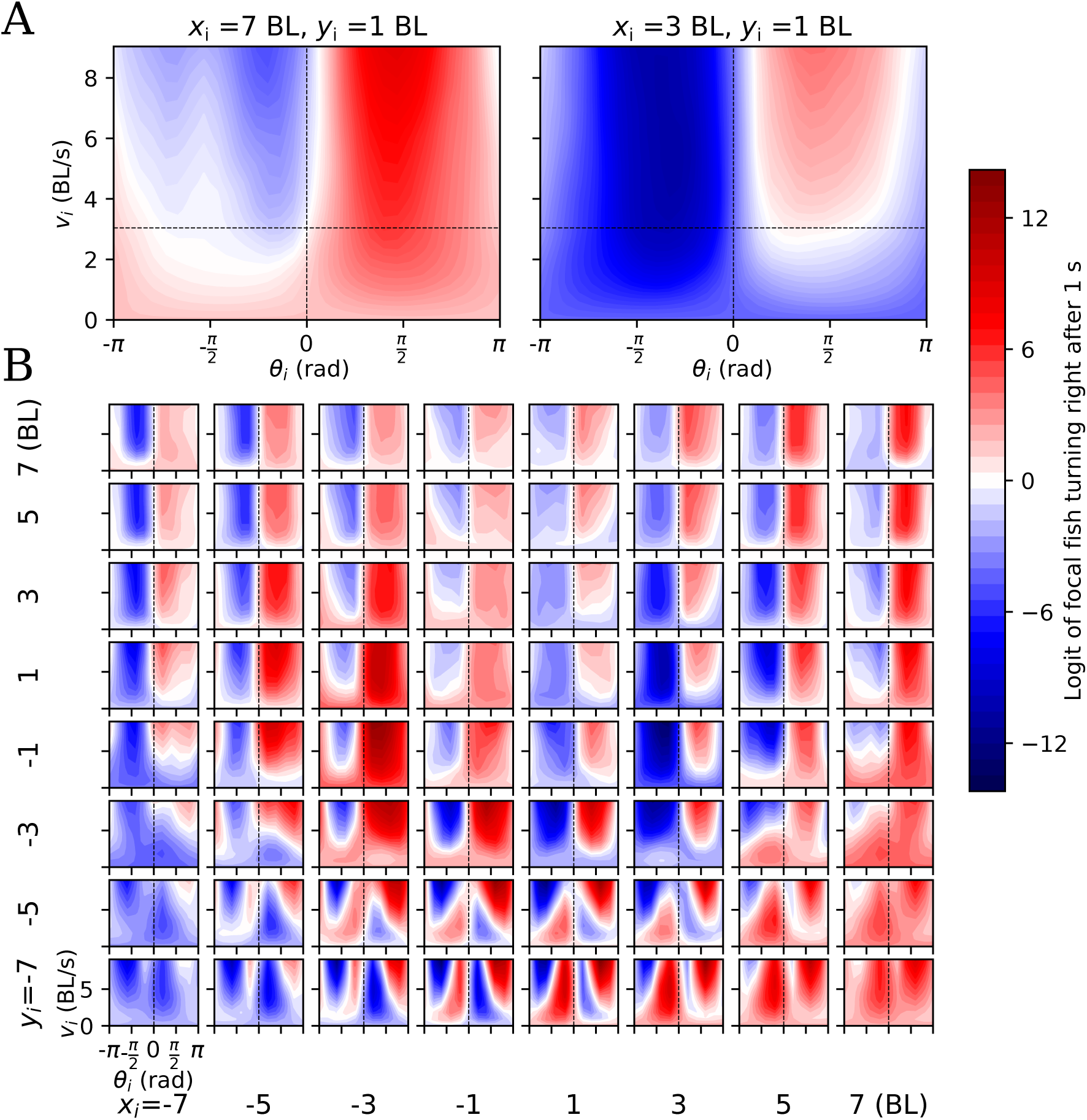
Properties of interaction between a pair of fish in the collective. (**A**) Logit *z* resulting from the pair-interaction subnetwork of the attention network, plotted as a function of the orientation of the neighbour respect to the focal, *θ_i_*, and speed of the neighbour, *v_i_*, for neighbour located at (*x_i_, y_i_*) = (7,1) BL (left) and (*x_i_, y_i_*) = (3, 1) BL (right). Focal speed is fixed at median velocity of 3.04 BL/s and focal acceleration at *a*_⊥_ =0 BL/s^2^. Red colour is evidence that the focal fish will turn right in 1s, while blue is evidence that the focal fish will turn left. Horizontal dashed line highlights the median speed of 3.04 BL/s. (B) Same as (A) but for 64 different neighbour positions (*x_i_, y_i_*), with *x_i_* and *y_i_* taking values in (−7, −5, −3, −1,1, 3, 5, 7) BL.

As a contrasting example, consider when the neighbour is closer and slightly in front, at *x_i_* = 3 BL and *y_i_* = 1 BL (Figure 2A, right). In this case, the focal gets repelled by the neighbour when the neighbor speed is below 3 BL/s.

These two examples illustrate how alignment, attraction and repulsion depend not only on the neighbour location but also on its speed (Figure 2B, similar to Figure 2A but for a 8×8 matrix of subplots, each for a different neighbour position), and on the speed (Figure S9, Figure S10, Figure S11, Figure S12) and acceleration of the focal (Figure S13).

From this six-dimensional function we can define alignment regions as those where the logit changes sign with neighbour orientation, that is, when focal will turn right (left) if neighbour orients to the right (left) (Figure 3, gray regions). The alignment score (Equation 11) measures how sensitive the logit is to neighbour orientation (Figure 3A, gray region; focal speed fixed at median value of 3.04 BL/s and neighbour speed indicated on top of each subplot). The alignment region increases in size and in score with increasing neighbour speed. At high neighbour velocities, strong alignment areas are 2-5 BL behind the focal and 3-5 BL at the sides (Figure 3A, right, darker gray regions). In a region 5-7 BL behind the focal there is a weak orientation effect but reversed in sign, with focal turning right (left) when neighbour orients to the left (right), (Figure 3A, pink). This anti-alignment region extends when increasing focal speed, while keeping neighbour speed fixed at the median value of 3.04 BL/s (Figure 3B, pink).

**Figure 3:**
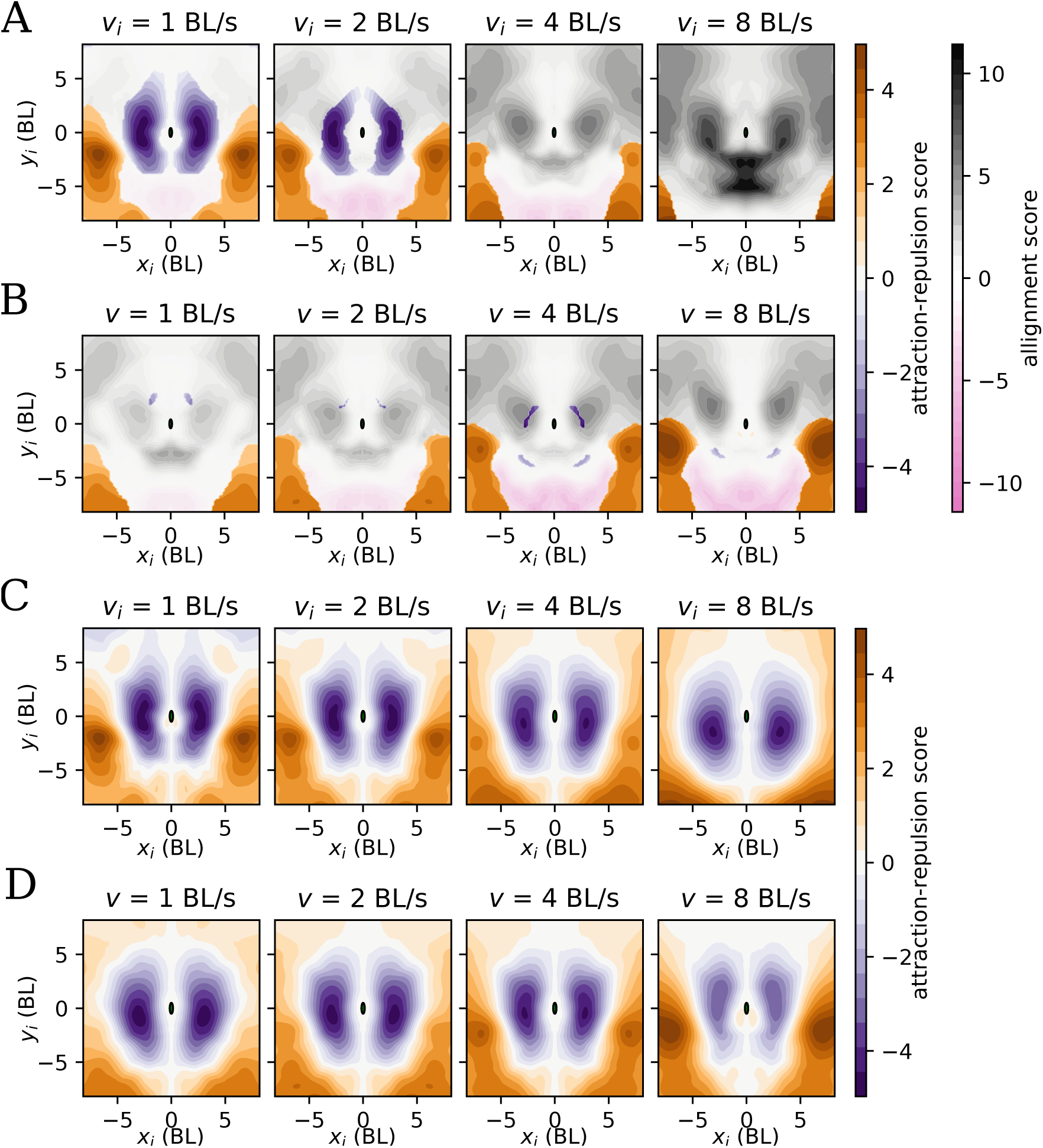
Alignment, attraction and repulsion zones depend on kinematic parameters of focal and neighbour. (**A, B**). Alignment (gray), attraction (orange), repulsion (purple) and anti-alignment (pink) zones. Alignment score (gray) measures how much the the logit changes when changing the neighbour orientation angle, and it is computed for those regions where there is an alignment effect defined by a change in sign of the logit when changing neighbour orientation angle. Attraction (orange) and repulsion scores (purple) are the logit averaged across relative orientation angles (positive or negative, respectively), plotted for regions with no alignment effect. (**A**) Scores given at four different values of the neighbour speed (1, 2, 4 and 8 BL/s) while fixing focal speed at the median 3.04 BL/s. Focal normal velocity fixed at *a*_⊥_ = 0. (**B**) Same as (**A**) but now fixing neighbour speed and varying focal speed. (**C, D**) Attraction and repulsion scores as in (**A,B**) but now plotted for all regions regardless of whether there is alignment effect or not.

We define attraction (resp. repulsion) regions as those where the logit does not change sign when changing the neighbour angle. Instead, the focal is attracted towards (resp. repelled from) the neighbour’s location independently of its orientation. The attraction-repulsion score (Equation 10) measures how positive (attraction) or negative (repulsion) is the logit of turning towards the neighbour (Figure 3A). Attraction regions shrink with increasing neighbour speed. They are mainly located to the side at 6-8 BL, extending to the back. Repulsion takes place only when the neighbour speed is below the median speed and neighbours are close to the focal (Figure 3A,B, purple).

However, classifying interactions into only 4 classes is oversimplistic, and a more complete account is captured by the six dimensional pair-interaction function in Figure 2. For example, when the neighbour is at (*x_i_*, *y_i_*) = (3, 1) BL and at high velocity, there is alignment but with a much higher probability of turning left at angles below *π*/2 than turning right at angles above *π*/2. This asymmetry in angles makes the sensitivity to orientation to the neighbour a mix of alignment and repulsion. We can see the full extent of relative attraction and repulsion zones by plotting the attraction-repulsion score for all points in space regardless of whether they correspond to an alignment effect or not (Figure 3C for different neighbour speeds and Figure 3D for different focal speeds). There is an approximately 5 BL diameter region of relative repulsion around the focal. Regions with a mix of alignment and repulsion (or attraction) are those of alignment in Figure 3A,B that overlap with regions of relative repulsion (attraction) in Figure 3C,D.

### How information is aggregated

The aggregation subnetwork *W* in Equation 1 outputs the (positive) weight of each neighbor in the aggregation. We found it to depend mainly on 4 variables

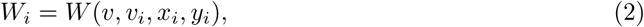

with *v* the focal speed, *v_i_* the neighbor speed and *x_i_* and *y_i_* the relative position of neighbour *i*.

In each subplot of Figure 4 we give *W* for different neighbour positions, keeping neighbour and focal speed constant. Generally, *W* is higher for neighbours that are closer to the focal, and lower for neighbours behind the focal. In the upper row of Figure 4 all subplots have the same focal speed at the median velocity of 3.04 BL/s, and each indicates the neighbour speed on top, with values *v_i_* = 1,2,4 and 8 BL/s. We see how *W* increases with neighbour speed for most neighbour positions, implying that faster neighbours carry more weight in the aggregation. This is more pronounced when the neighbour is close by and to the side.

**Figure 4:**
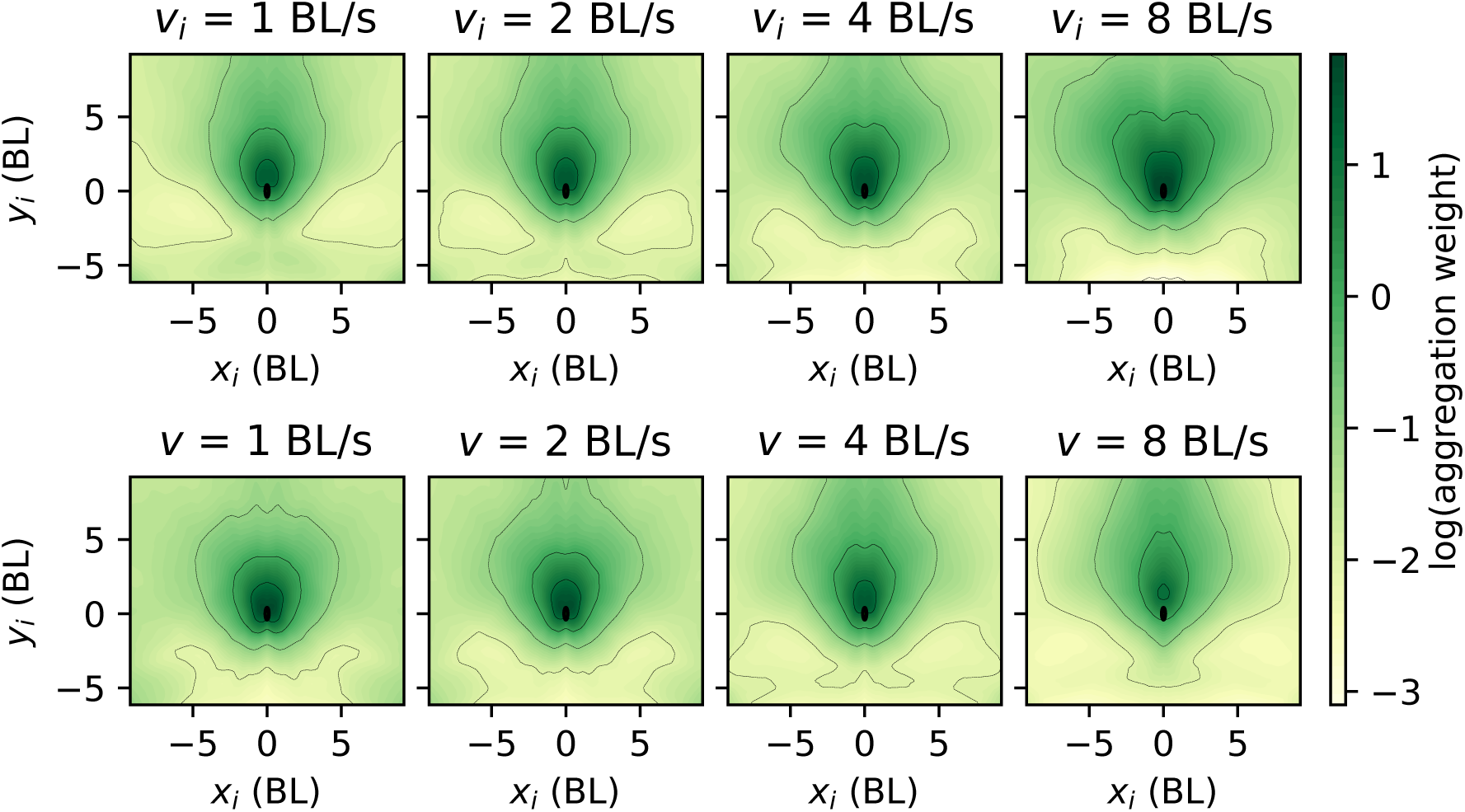
Weighting function: How a fish aggregates information from neighbours. Logarithm of the aggregation weight, log(*W*), as a function of neighbour position, *x_i_* and *y_i_*. Top row: focal speed fixed at 3.04 BL/s and each subplot corresponding to different neighbour speeds marked on top of each. Bottom row: same as top row but for fixed neighbour speed at 3.04 BL/s and different focal speeds.

In the lower row of Figure 4 all subplots have the same neighbour speed at the median velocity of 3.04 BL/s, and each indicates the focal speed on top, with values *v_i_* = 1,2,4 and 8 BL/s. We see how the mass of *W* increasingly shifts towards the front the faster the focal fish moves. Adding other variables to the attention marginally improves accuracy, and still further insight is gained. When the neighbour orientation angle is added, higher values of the weight *W* are obtained in positions leading to an immediate collision (Figure S14).

The final impact of each neighbour on the probability of the focal turning right can be seen from Equation 1 to be given by a normalized weight, that is, the weight given by *W* relative to the sum of the weights of all individuals,

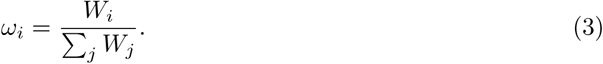

For example, if all 25 neighbours are assigned the same weight by *W*, after normalization all animals weight 1/25 = 0.04, no matter how large or small the value of *W* is. If one of the neighbours has a higher (lower) value of *W*, the importance of the other neighbours decreases (increases).

We give the normalized weights *ω_i_* in Equation 3 for each neighbour in three illustrative frames (Figure 5A; animals with higher normalized weights in darker green). We observe cases in which a large number of neighbours has an important impact (Figure 5A, lower right), others with fewer (Figure 5A, upper right) or even with a single neighbour that overweighs the others by far (Figure 5A, upper left).

**Figure 5:**
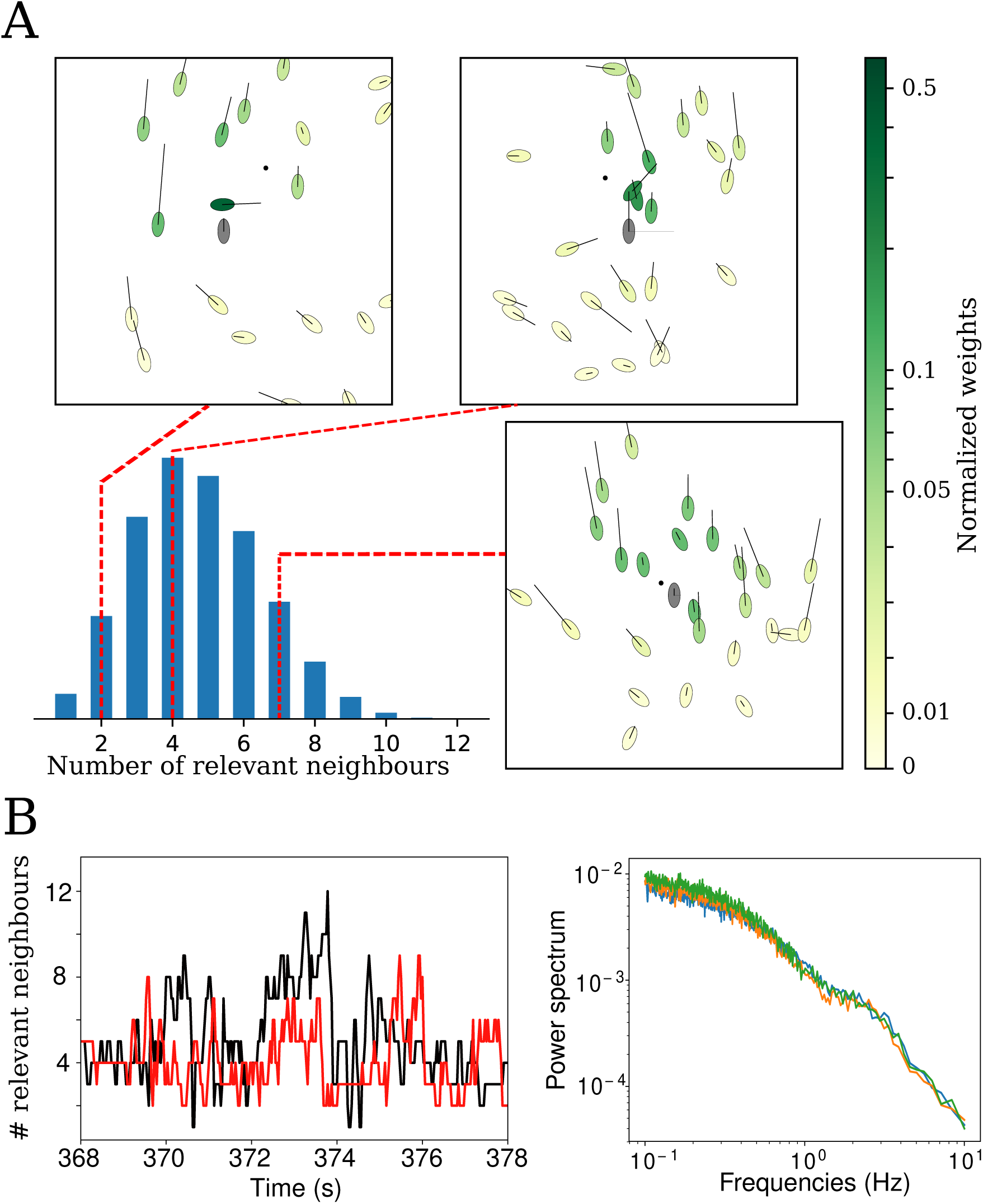
Relevant neighbours in the aggregation. **A**. Distribution of number of relevant neighbours. Three example frames with each neighbour coloured with its normalized weight in the aggregation. Focal animal is indicated in gray color, with a horizontal line proportional to the normal acceleration to either left or right and a small dot in its frontal positions indicating the focal position 1 second into the future. In both focal and neighbours, a line along the major axis indicates the fish velocity. **B** Left: Time variation of the number of relevant neighbours for two focal fish and for a short period of time. Right: Power spectrum of the number of relevant neighbours. Colours correspond to 3 different videos.

To quantify the number of relevant neighbours in each frame, we extracted those with a weight larger or equal to the typical one in the frame, *ω*_t_ = exp (∑*_i_ ω_i_* log(*ω_i_*)) ([27], section 4.4). When all individuals have the same weight, this criterion gives the total number of neighbours in the model, 25. In the case of one neighbour outweighting all the others by far, it can can give 1 as a result. The resulting distribution of relevant neighbours has a mode of 4 neighbours, a mean of 4.7, a standard deviation of 1.8 (Figure 5A). To gain some intuition on the dynamics of the number of relevant neigbours, consider how this value changes in time for two focal individuals (Figure 5B, left). In these two examples, it can be seen that the focal has often 4 relevant neighbours but this number fluctuates and can shift to higher and lower numbers in a fast subsecond scale. We analyzed the time scales by computing the power spectrum (Figure 5B, right; shown for 3 videos of 100 animals each). The power spectrum monotonically decreases with frequency, implying that most of the variation occurs at low frequencies, albeit without a clear time-scale (Figure 5B, right)

## Discussion

Our results show that animals in collectives can use an aggregation rule that naturally allows individuals to shift from simple local averaging to following a single individual. This rule may be seen as allowing a smooth transition from average type models [3, 4, 5, 6, 28, 7, 20] and models in which one or very few animals influence the rest as in the many-eyes model for predator detection [8, 9, 10, 29, 11] and others [17]. Note that this ability to shift from many to few can allow a collective to match the changing knowledge distribution in the group [30]. We believe our methodology could open the door to the experimental study of the properties of this matching.

From the pairwise interaction in the collective, we could extract attraction, repulsion and alignment as approximate notions [4, 6]. Usually, these interaction classes are defined only in terms of relative position of neighbour. However, we found them to exist in a 6-dimensional space. This translates into these classes also depending on speeds of focal and neighbour, focal acceleration and relative orientation between the two fish. This implies that experiments testing for the relevance of one variable, say speed, may give contradictory results depending on the analysis strategy. Our results imply that analysis needs to take into account that the interactions take place in a space with more dimensions. Also note that the three classes are not cleanly separated as alignment regions are mixed with attraction or repulsion, as found in [31].

As a strategy to extract the relevant variables for behavior, we have required them to predict future behavior (like e.g. [32, 33, 25, 20]). This approach has the additional advantage of automatically generating labelled data for supervised training of networks. It can be enriched, at the cost of increasing the model dimensionality, with more information about behavioral history, possible internal variables (parametrized, for example, by time of day or more direct internal measurements), explicit dynamics and posture using reduced variables [34, 35, 36, 37]. A second requirement for our models was that they should work for data not used to obtain the model. These two requirements are standard in machine learning, but less so in the biological sciences.

Our results illustrate how modular deep networks enable flexible data-driven modelling without losing insight. Each module is flexible, with tens of thousands of parameters, but implements a function with low dimensionality in the number of inputs and outputs. Combinations of modules [38, 39], two types in the attention network considered, achieve higher compositional complexity that adds flexibility without losing insight.

## Methods and Materials

### Data and code availability

60- and 100-fish as well as the new 80-fish videos can be found at www.idtracker.ai. Code used in this study is free and open-source and may be used to study interactions in any animal species or other agents (https://gitlab.com/polavieja_lab/fishandra).

### Animal rearing and handling

Zebrafish, *D. rerio*, of the wild-type TU strain were raised by the Champalimaud Foundation Fish Platform, according to methods in [40]. Experimental procedures were approved by the Champalimaud Foundation Ethics Committee and the Portuguese Direcção Geral Veterinária in accordance to to the European Directive 2010/63/EU. Handling procedures were as in [21]. We used juveniles of 31-33 days post fertilization.

### Videos and tracking

We used 6 videos of 60 and 100 freely swimming juvenile zebrafish from [21], and 3 new videos of 80 juveniles. The camera had a frame rate of 32 fps and 20 Mpx of definition. We obtained all the fish trajectories using *idtracker.ai* with an accuracy of 99.95 (mean) ±0.01% (std) [21].

### Preprocessing

We interpolated linearly the very small holes in the tracked trajectories (0.027% for 100-fish videos). We normalized trajectories, by translation (center of arena at (0,0)) and scaling (radius of the arena at 1). To reduce noise while preventing contamination by any future information, we smoothed the trajectories using a 5-frame half-Gaussian kernel with *σ* = 1 frame. We obtained velocity and acceleration by finite differences, using only current and past frames. To avoid direct border effects, we removed datapoints where the focal fish is further away from the center than 80% of the radius. Each video was divided in three parts, to obtain the training, validation and test datasets (97%/2%/1%).

In each video frame, for each individual, we found the *n* nearest neighbours 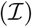. We then obtained (i) velocity and acceleration of the focal fish, (ii) relative position, absolute velocity and absolute acceleration of the closest *n* neighbours, (iii) whether the focal fish has turned right or left after *N_f_* frames in the future.

### Deep networks

We implemented the Deep Networks using Keras [41] through its Python API and with TensorFlow backend [42]. We solved the following classification task: Given dynamical properties of a focal fish and its *n* closest neighbours, does the focal fish turn right or left after 1s? Asocial information is the set of speed and normal and tangential acceleration of the focal,

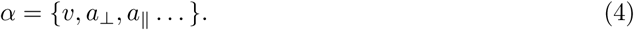

Social information from a neighbour *i* is its location, velocity and acceleration

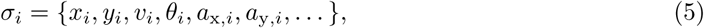

whose coordinates we calculate in an instantaneous frame of reference that is not moving, which is centered in the focal fish and whose y-axis is co-lineal with the focal fish velocity. Note that *v_i_* is the absolute speed, while (*x_i_*, *y_i_*) is the relative position of the neighbour, rotated to the frame of reference. In each network, we first obtain the logits *z*, and then the probabilities by using a logistic function *p* = 1/(1 + *e*^−^*^z^*).

### Interaction network

In the interaction network [26], given asocial (*α*) and social 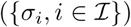 information, the logit of turning right is calculated as

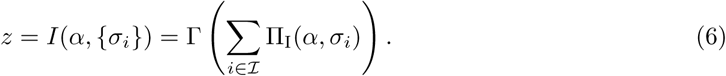

The function Π_I_ is the pair-interaction subnetwork. We modelled it using a fully-connected network with 3 hidden layers of 128 neurons each, plus a readout layer of 128 neurons. There are rectified linear unit (ReLU, [43]) nonlinearities after each hidden layer (but not after the readout). The outputs of Π_I_ for different neighbours are summed together and transformed by a second function, Γ. We modelled Γ as a fully-connected layer with one hidden layer of 128 neurons, plus a one-neuron readout layer. There are ReLU nonlinearities preceding the whole network and after each hidden layer (but not after the one-neuron readout layer).

To effectively multiply available data by *n*, we considered all neighbours to be equal. Equivalently, there is symmetry with respect to exchange of neighbour labels. We did not observe any turning side preference. Therefore, to effectively multiply available data by 2, we forced the network to be antisymmetrical with respect to a reflection along the body axis by antisymmetrization of *I*,

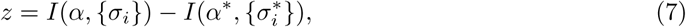

where the star superscript represents a reflection along the longitudinal axis of the body, calculated by switching the sign of all *x* components.

### Attention network

Equation 1 can be rewritten using a notation that compares directly with Equation 6 as

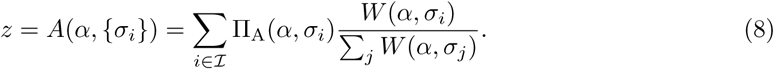

The function Π_Α_ captures the effect of pairwise interactions. It has the same structure as Π_I_ except that its readout layer has only one neuron, and that we antisymmetrise it. *W* is an attention layer, weighting the logits of the different neighbours. *W* has the same structure as Π_Α_, except that it accepts as input a y-axis-reflection-invariant subset of the asocial and social variables, and that there is an exponential function after the single-neuron readout signal.

### Loss

Following standard procedures in binary classification, when training the network to estimate the probability *p_i_* of turning right, we minimised the cross-entropy loss, [43]

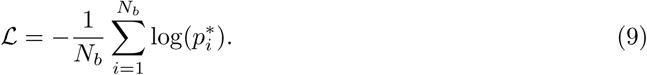

summed along *N_b_* data points in the minibatch and where 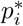 is the probability given by the network to the actual turn. When the network predicts a right turn with probability *p_i_*, 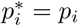 if the actual turn was to the right, and 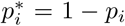 if the actual turn was to the left. We minimise loss using Adam [43]. We stopped training if validation loss did not reach a new minimum for 10 epochs and did increase 25% from the current minimum, or after 100 training epochs. In the attention network, we annealed learning rate from 10^−4^ to 10^−5^, using a batch size of 500. In the attention network, we annealed learning rate from 5 × 10^−5^ to 10^−5^ and trained with a batch size of 200. Dropout [43] did not improve accuracy.

### Attraction-repulsion and alignment scores

Attraction-repulsion score is obtained as

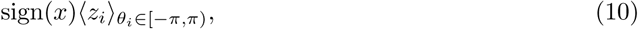

where 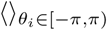 indicates average over all possible relative orientation angles of the neighbour. There is attraction (repulsion) when the score is positive (negative). Alignment score is obtained as

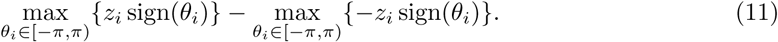

## Acknowledgments

We thank Kristin Branson, Jimmy Liao and the Collective Behavior Lab at Champalimaud for feedback and Ana Catarina Certal for animal care. This work was supported by Fundação para a Ciência e Tecnología PTDC/NEU-SCC/0948/2014 (to GGdP and contract to FJHH), Congento LISBOA-01-0145-FEDER-022170, NVIDIA (FJHH, GGdP), Champalimaud Foundation (GGdP) and GGdP work at KITP Santa Barbara by NSF Grant No. PHY-1748958, NIH Grant No. R25GM067110, and the Gordon and Betty Moore Foundation Grant No. 2919.01. F. R-F. acknowledges a FCT PhD fellowship.

---

**Table S1:** Accuracy of prediction of the turning side of the focal fish after 1 second evaluated on a held-out test data of large turns (20°-160°, test data) when using different variables. 25 neighbours, interaction network, model with lowest validation loss of two runs. In all cases, in addition to the variables mentioned in the table, we provide the relative position of the neighbour (*x_i_*, *y_i_*). Elsewhere in this article, we use *v*, *a*_⊥_, (*v_i,x,_ v_i,y_*) and (*x_i,_ y_i_*), the simplest among the sets of variables with high accuracy.

|  |  | Neighbour |  |  |
| --- | --- | --- | --- | --- |
| | | $(a_{i,x}, a_{i,y})$ | $(v_{i,x}, v_{i,y})$ | $(v_{i,x}, v_{i,y})$ & $(a_{i,x}, a_{i,y})$ |
| Focal | $(a_{\perp}, a_{\parallel})$ | 81.6% | 84.1% | 84.2% |
| | $v$ | 80.7% | 83.9% | 83.7% |
| | $v$ & $(a_{\perp}, a_{\parallel})$ | 81.5% | **84.5%** | **84.5%** |
| | $v$ & $a_{\parallel}$ | 80.9% | 83.7% | 83.8% |
| | $v$ & $a_{\perp}$ | 81.5% | **84.5%** | **84.4%** |

**Table S2:** Different architectures of interaction networks (number neurons of per layer × number of hidden layers). Best (i.e. lowest validation loss) of at least three runs with different batch sizes.

| layers | layers | validation | Test accuracy |  |  |
| --- | --- | --- | --- | --- | --- |
| pair-wise | interaction | loss | all turns | 20°-160° | 30°-100° |
| 32 × 2 | 128 × 1 | 0.53359 | 74.2% | 83.8% | 86.2% |
| 32 × 3 | 128 × 1 | 0.52641 | 74.6% | 84.4% | 86.9% |
| 32 × 4 | 128 × 1 | 0.52774 | 74.5% | 84.2% | 86.5% |
| 32 × 5 | 128 × 1 | 0.52787 | 74.8% | 84.5% | 86.6% |
| 64 × 2 | 128 × 1 | 0.53207 | 74.2% | 83.8% | 86.0% |
| 64 × 3 | 128 × 1 | 0.52814 | 74.6% | 84.2% | 86.7% |
| 64 × 4 | 128 × 1 | 0.52785 | 74.7% | 84.5% | 86.8% |
| 64 × 5 | 128 × 1 | 0.52650 | 74.7% | 84.1% | 86.2% |
| 128 × 2 | 128 × 1 | 0.53238 | 74.5% | 84.2% | 86.7% |
| 128 × 3 | 128 × 1 | 0.52680 | 74.5% | 84.0% | 86.4% |
| 128 × 4 | 128 × 1 | **0.52559** | 74.6% | 84.5% | 87.1% |
| 128 × 5 | 128 × 1 | 0.52718 | 74.4% | 84.1% | 86.4% |
| 256 × 2 | 128 × 1 | 0.53157 | 74.5% | 84.0% | 86.5% |
| 256 × 3 | 128 × 1 | 0.52735 | 74.7% | 84.4% | 86.8% |
| 256 × 4 | 128 × 1 | 0.52740 | 74.8% | 84.5% | 87.0% |
| 256 × 5 | 128 × 1 | 0.52641 | 74.6% | 84.0% | 86.3% |

**Table S3:**
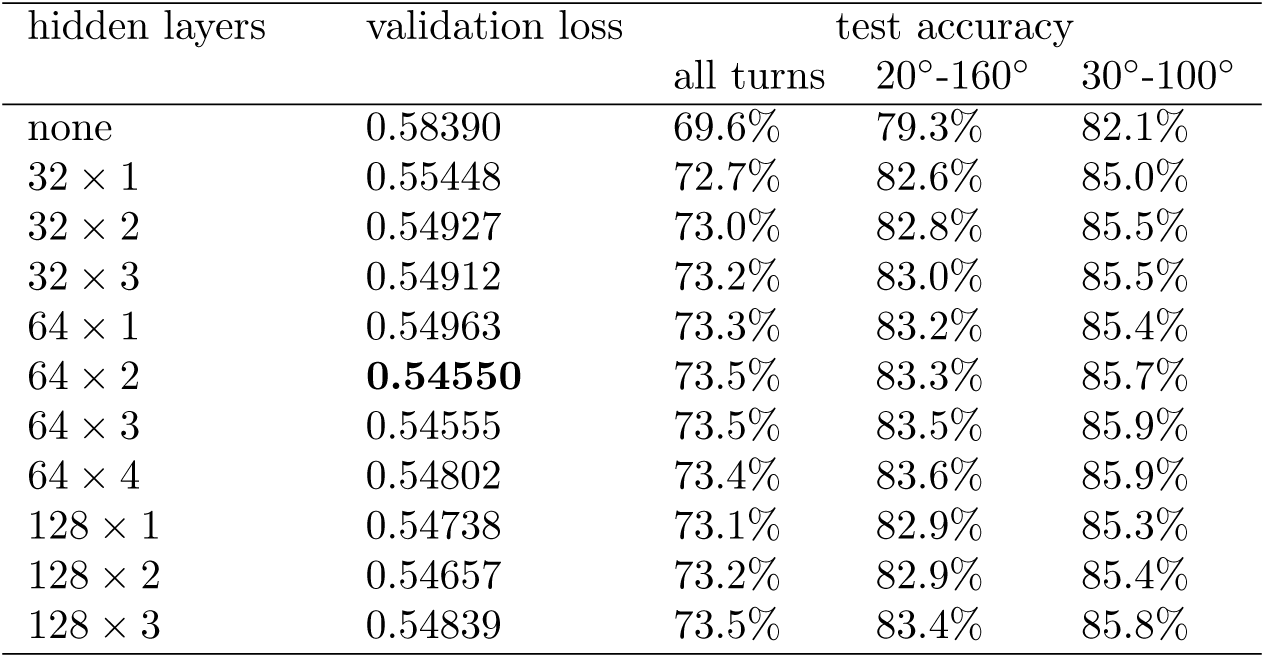
Different architectures of fully connected networks (number of neurons per layer × number of hidden layers). Best of at least three runs with different batch sizes.

**Figure S1:**
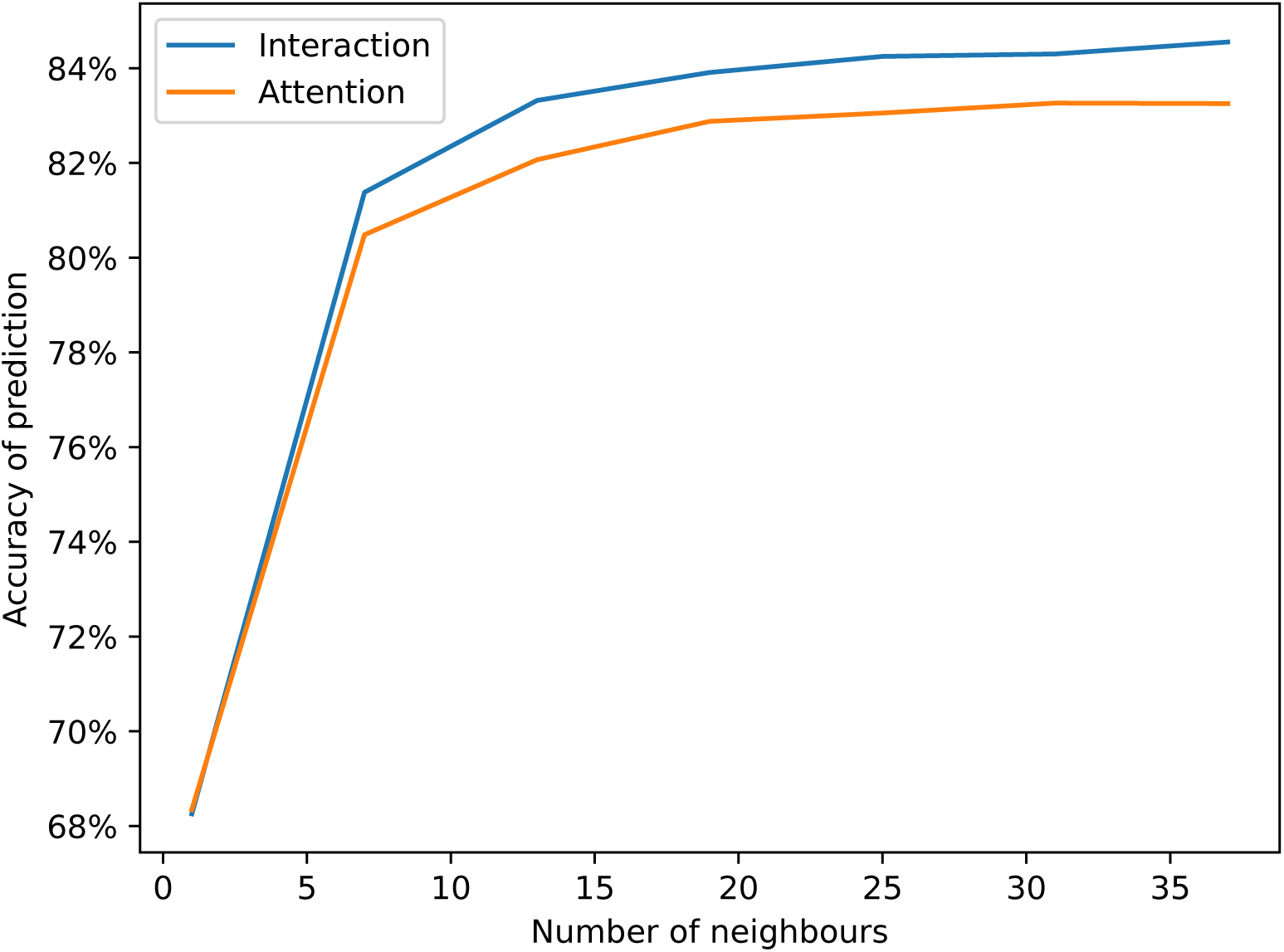
Accuracy of prediction of the turning side of the focal fish after 1 second evaluated on a held-out test data of large turns (20°-160°) as a function of number of closest neighbours. Both the interaction network (blue) and the attention network (orange) improve in accuracy with the number of neighbours, and then plateau after approx. 20 neighbours.

**Table S4:** Accuracy of the prediction of large turns for videos of different number of animals. 25 neighbours, model with lowest validation loss of two runs.

|  | Interaction | Attention |
| --- | --- | --- |
| 100 individuals | 84.4% | 83.2% |
| 80 individuals | 77.1% | 76.4% |
| 60 individuals | 73.4% | 72.2% |

**Figure S2:**
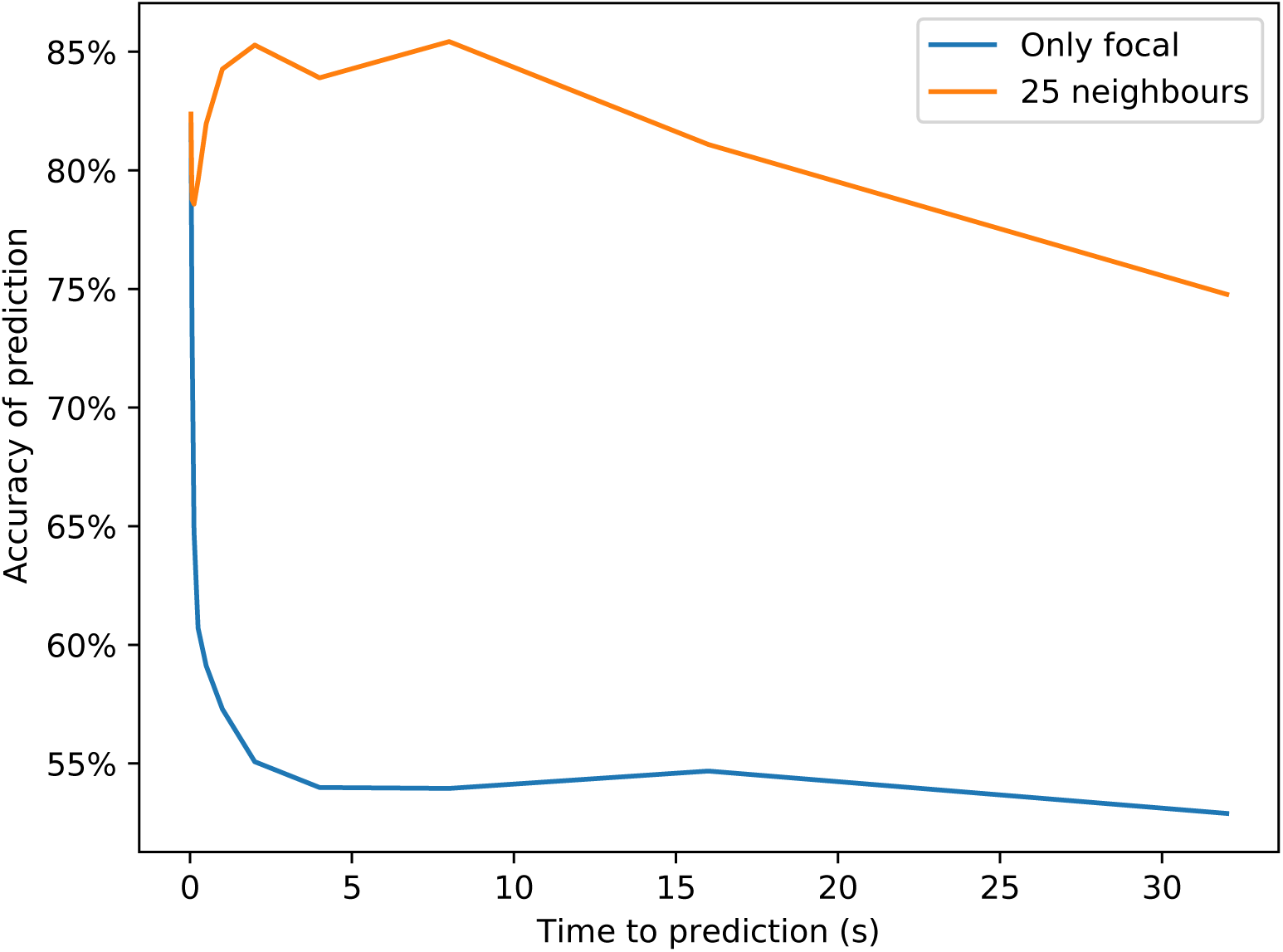
Accuracy of prediction of the turning side of the focal fish evaluated on held-out test data of large turns (20°-160°) for different times to prediction. The prediction from an interaction model with 25 neighbours (orange) and from a model that is blind to any social information (blue). Accuracy for immediate futures (<100 ms) is high for both models, because of correlations in the acceleration. Accuracy with 25 neighbours has a local minimum for futures of 250 ms, it has a broad maximum when predicting futures between 1 and 10 s, and then it drops slowly when predicting more distant futures.

**Figure S3:**
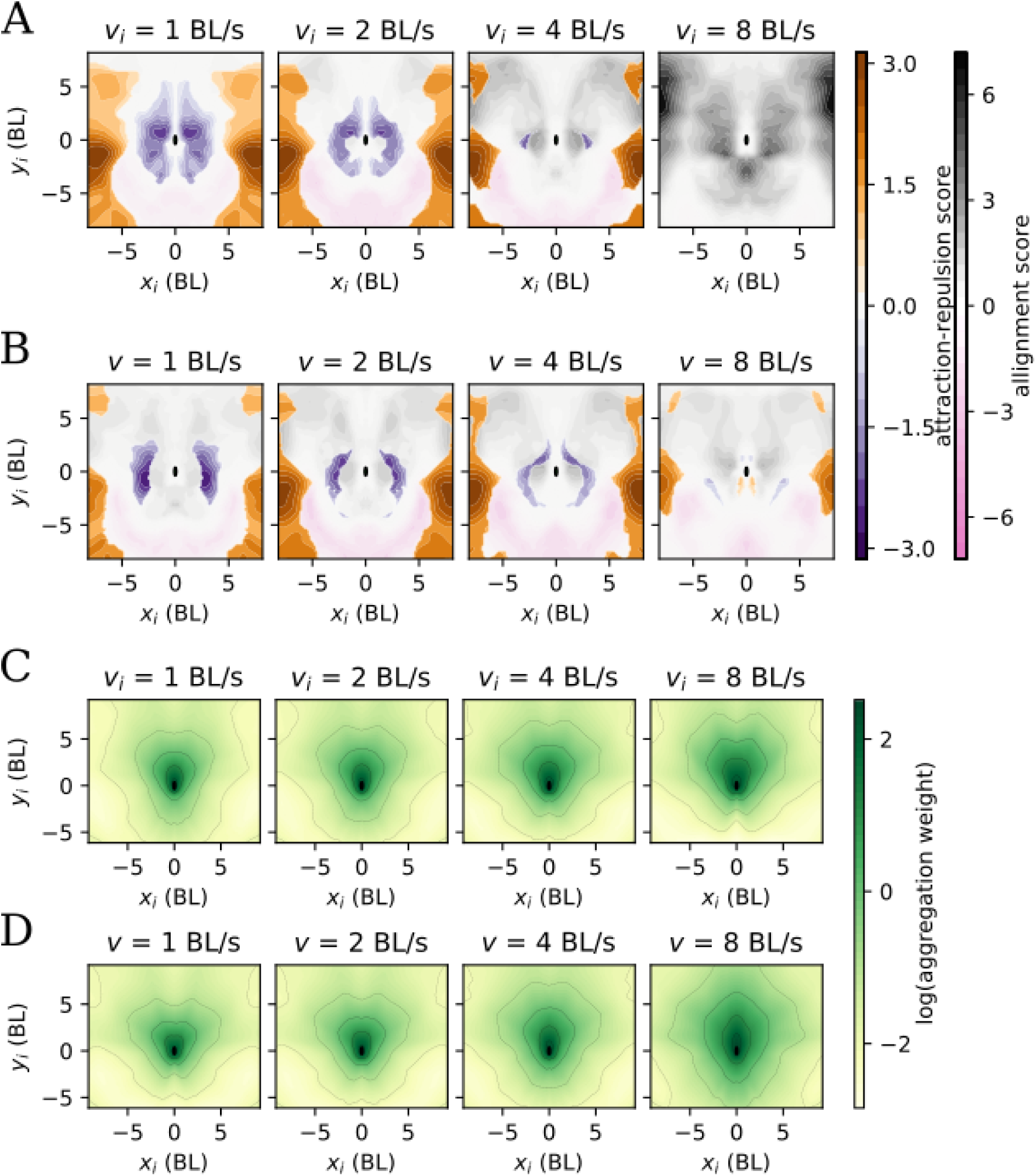
Pair interaction and aggregation in 250 ms predictions. **A** Same as Figure 3A. **B** Same as Figure 3B **C** Same as Figure 4. Note how high-attention areas are closer to the focal fish. **D** Same as Figure 4

**Figure S4:**
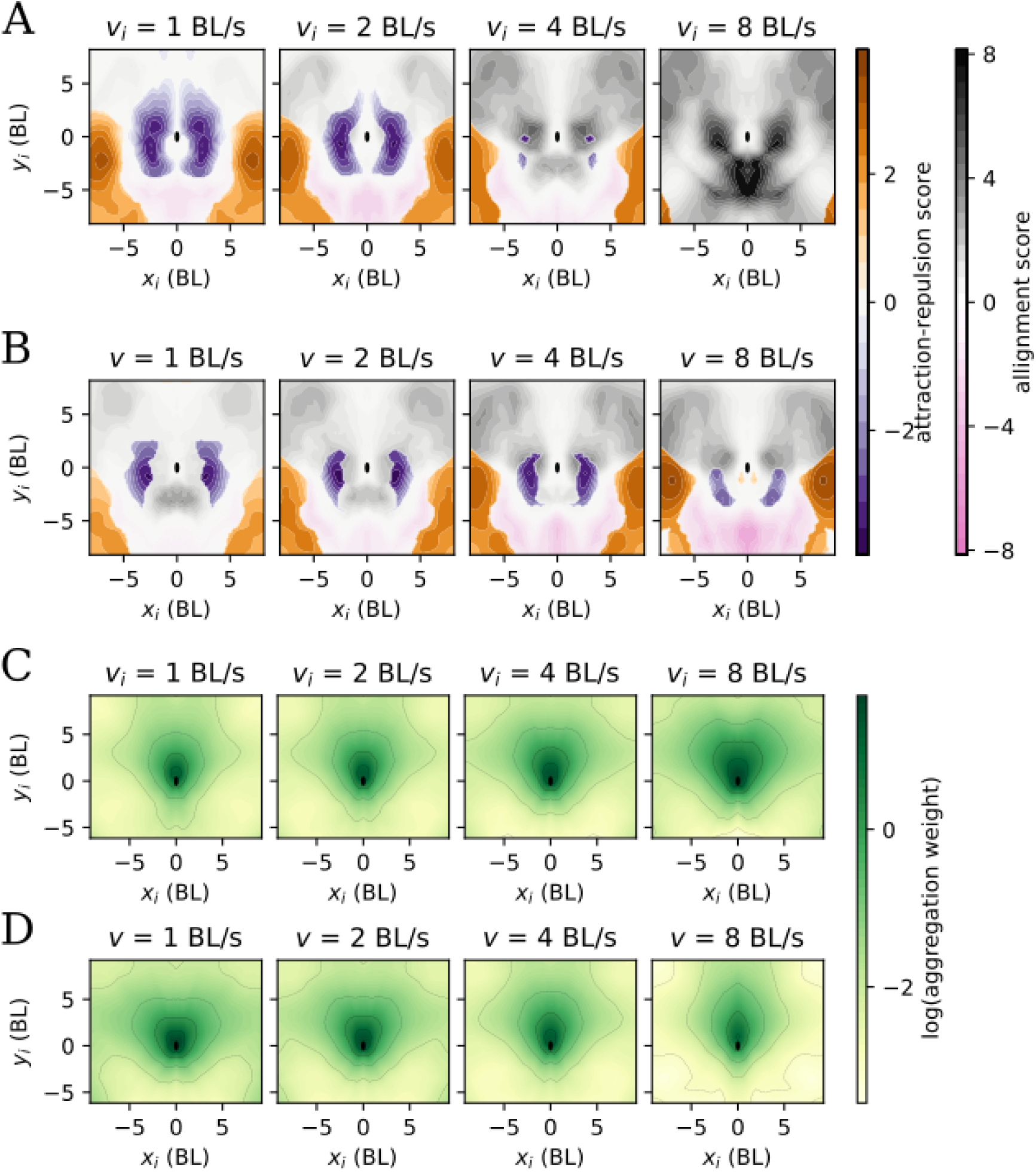
Pair interaction and aggregation in 500 ms predictions. **A** Same as Figure 3A. **B** Same as Figure 3B C Same as Figure 4. **D** Same as Figure 4

**Figure S5:**
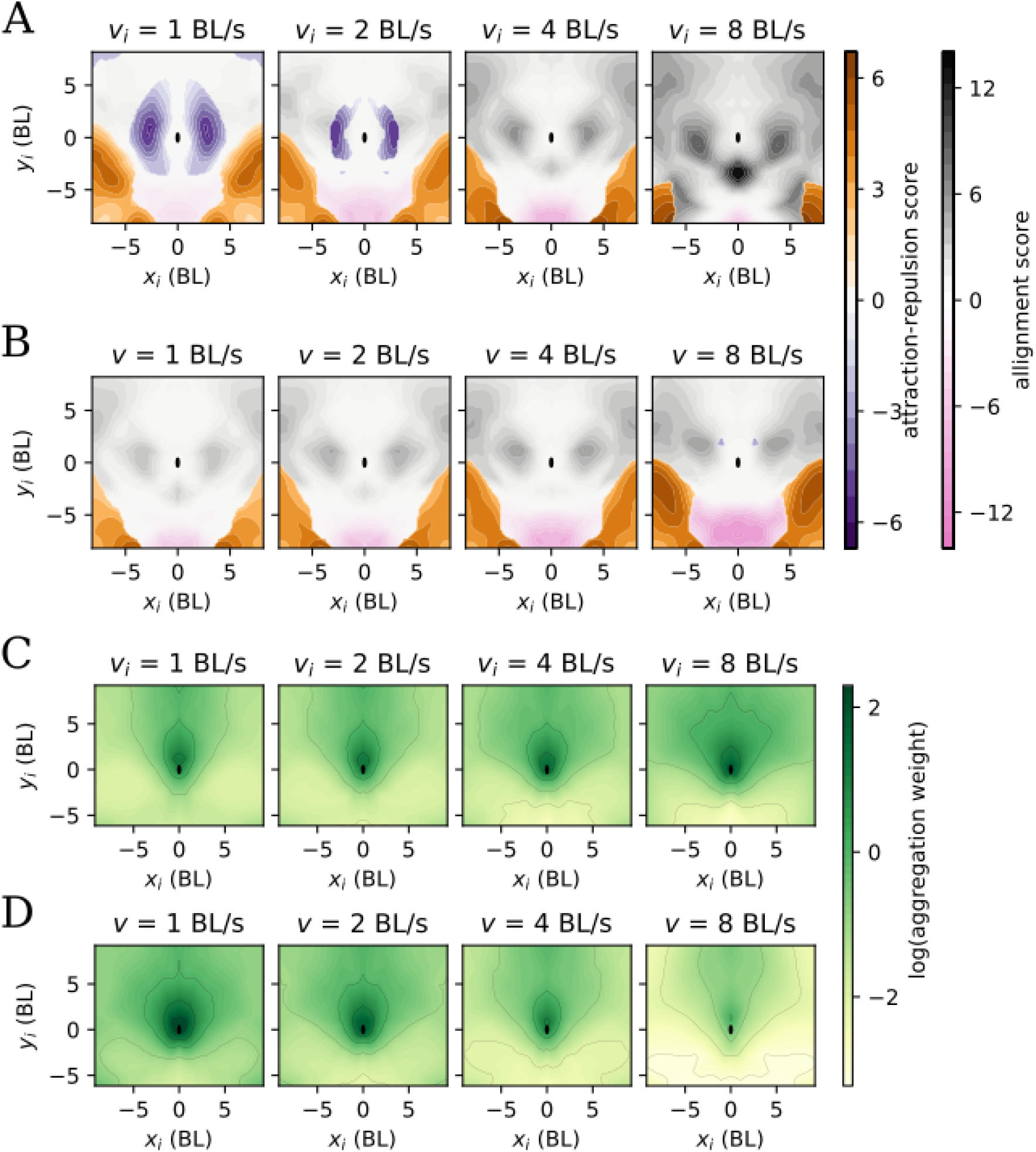
Pair interaction and aggregation in 1500 ms predictions. **A** Same as Figure 3A. **B** Same as Figure 3B **C** Same as Figure 4. Note how high-attention areas are closer to the front. **D** Same as Figure 4

**Figure S6:**
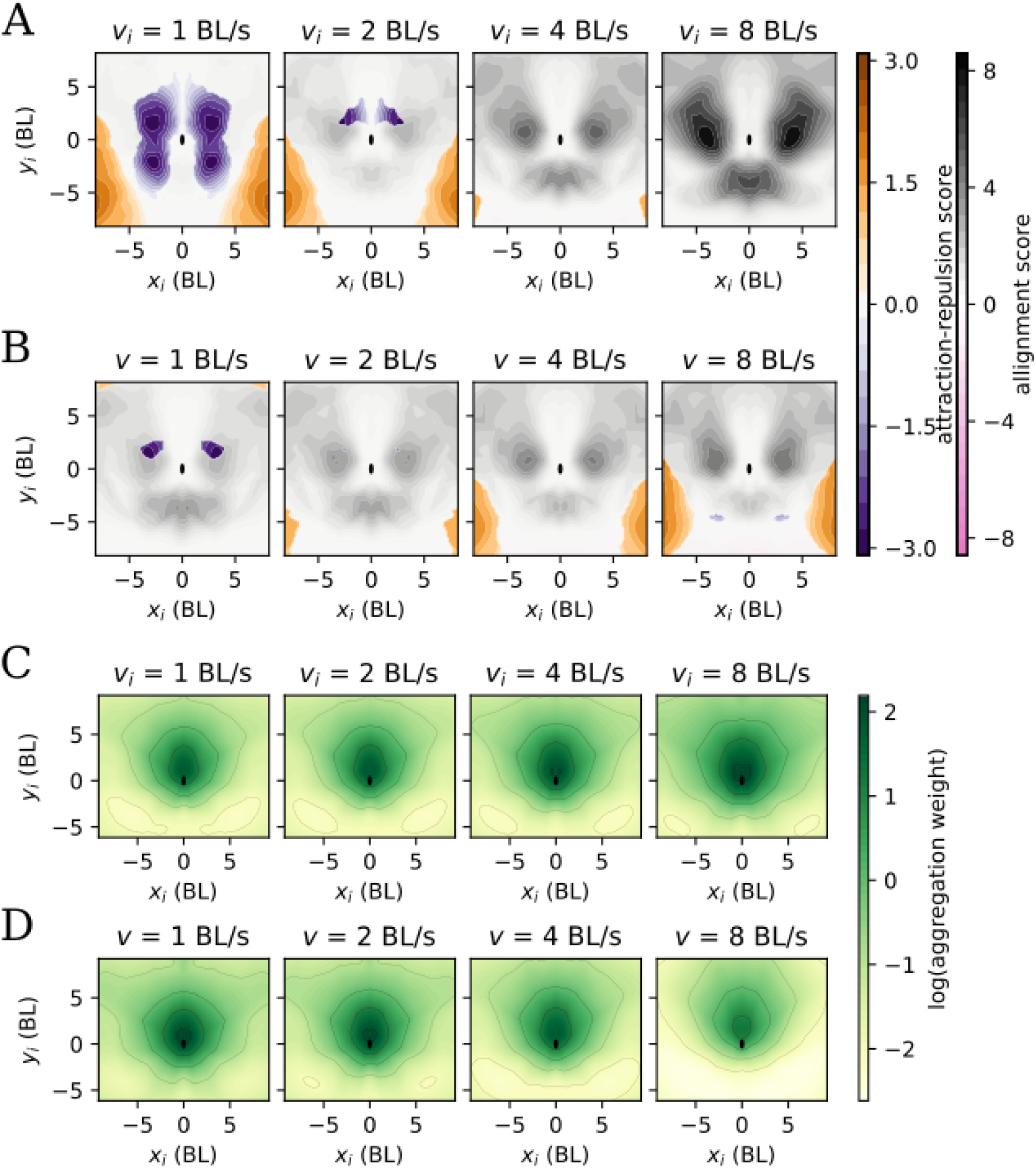
Pair interaction and aggregation, obtained from 60-fish videos. **A** Same as Figure 3A. The most conspicuous difference with Figure 3A is the weakening of anti-alignment **B** Same as Figure 3B **C** Same as Figure 4. **D** Same as Figure 4. Note the comparatively weak attention at the back of the focal.

**Figure S7:**
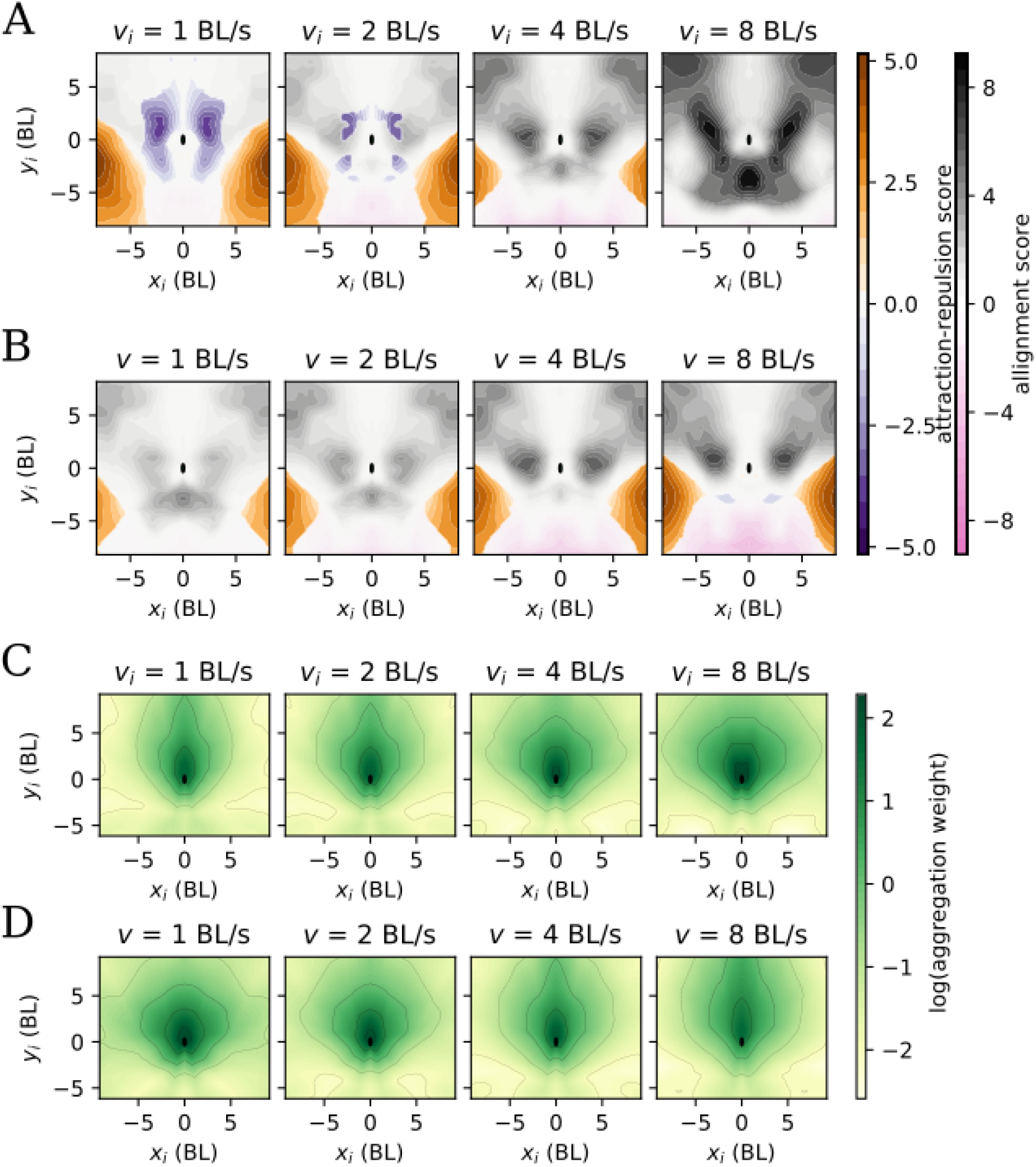
Pair interaction and aggregation, obtained from 80-fish videos. **A** Same as Figure 3A. The most conspicuous difference with Figure 3A is the weakening of anti-alignment **B** Same as Figure 3B **C** Same as Figure 4. **D** Same as Figure 4.

**Figure S8:**
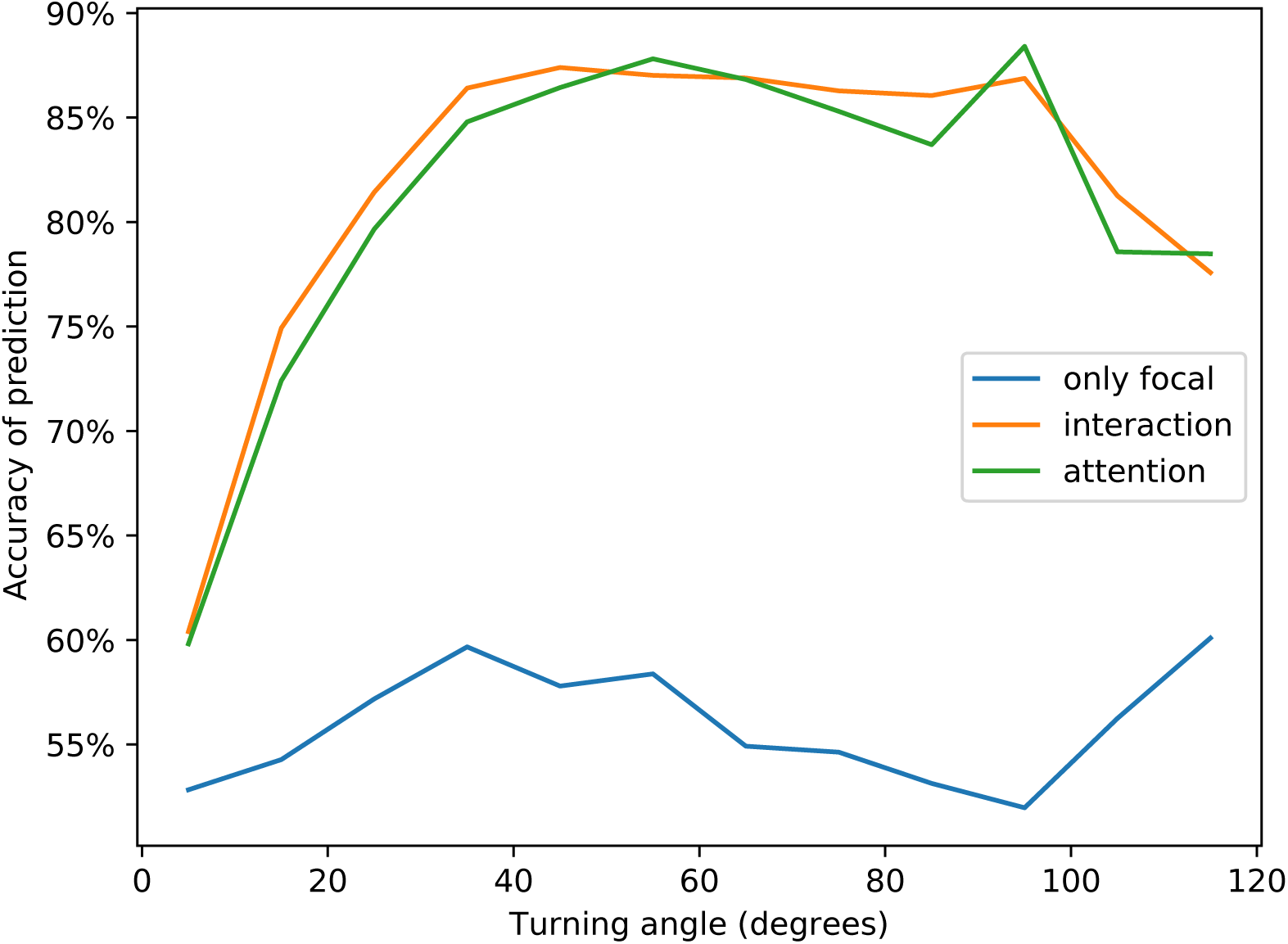
Accuracy of prediction of the turning side of the focal fish after 1 second evaluated on held-out test data of large turns (20°-160°). When using only focal variables (blue) remains low at all turning angles. Both networks integrating information from 25 neighbours, interaction (orange) and attention (green) perform better at turning angles between 40 and 100.

**Figure S9:**
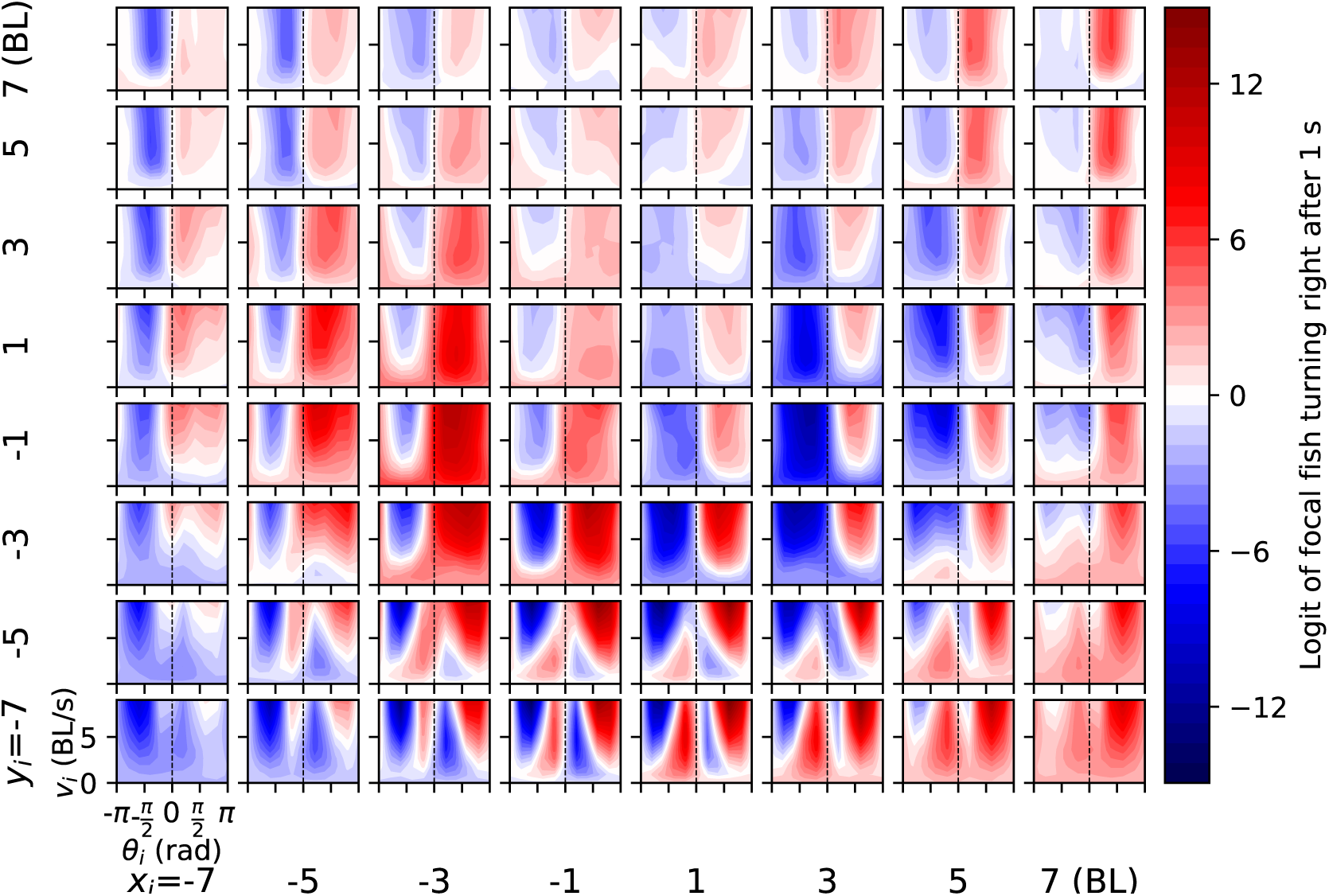
Properties of interaction between a pair of fish in the collective when the focal fish is moving at low speed (1 BL/s).

**Figure S10:**
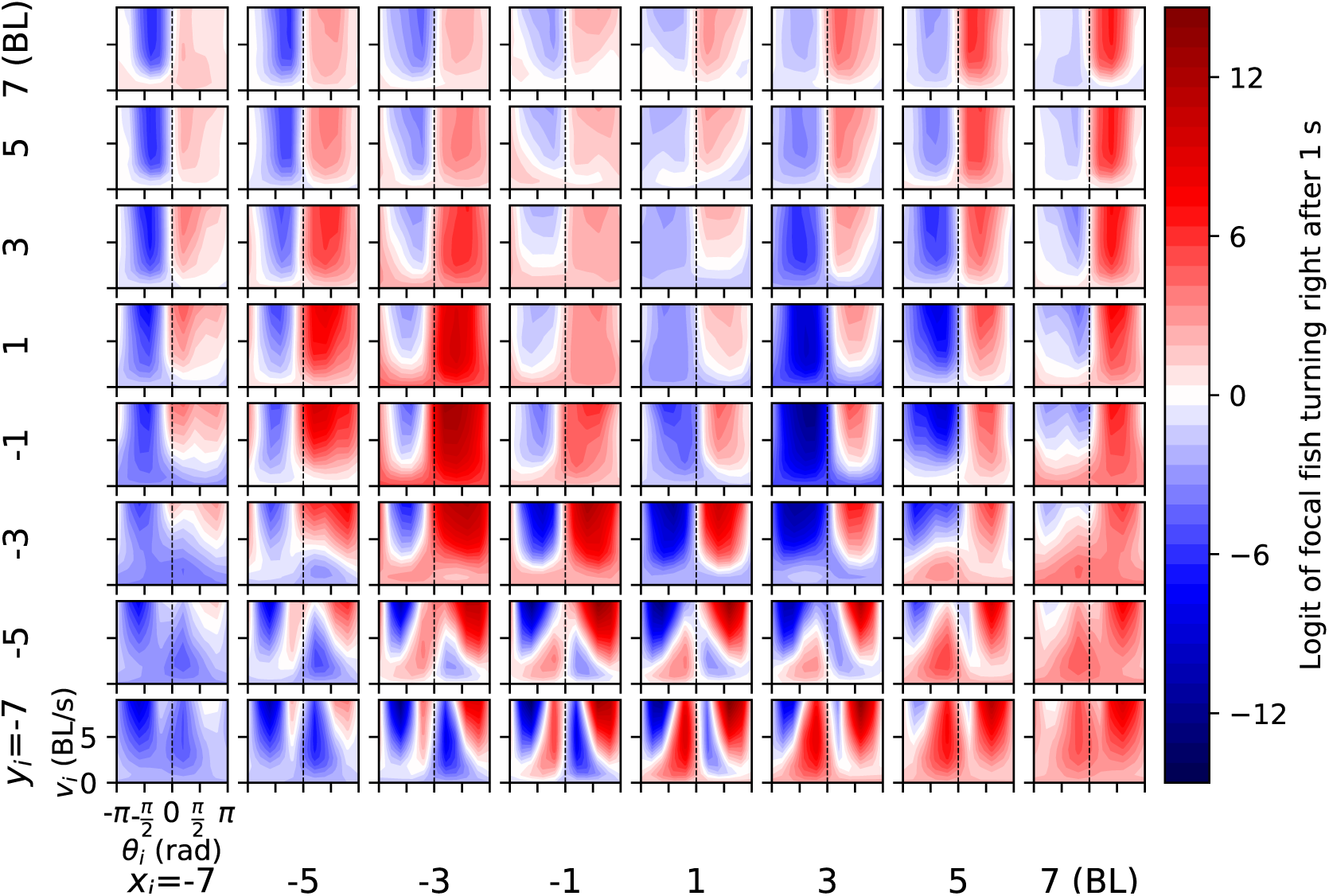
Properties of interaction between a pair of fish in the collective when the focal fish is moving at low speed (2 BL/s).

**Figure S11:**
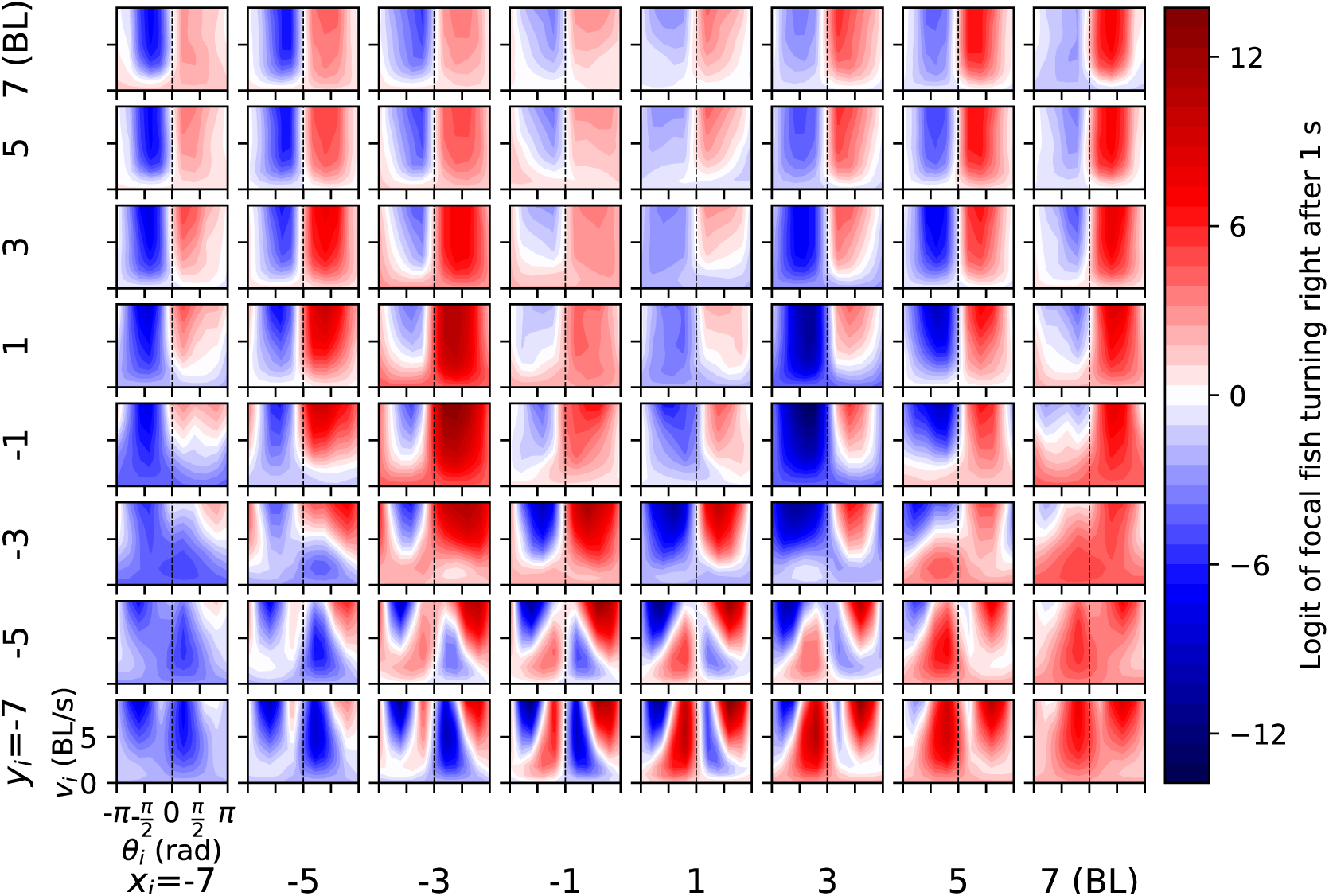
Properties of interaction between a pair of fish in the collective when the focal fish is moving at low speed (4 BL/s).

**Figure S12:**
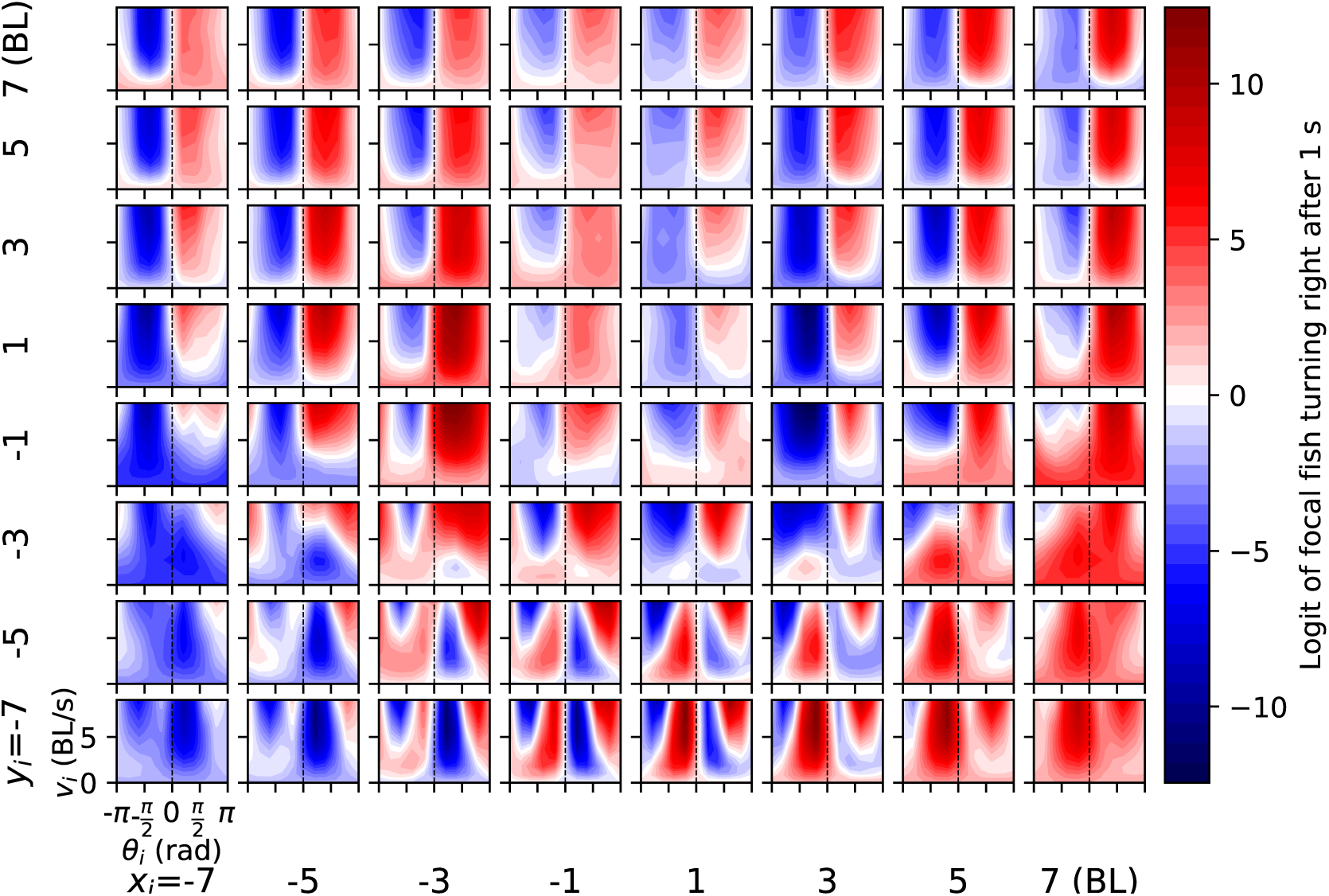
Properties of interaction between a pair of fish in the collective when the focal fish is moving at low speed (8 BL/s).

**Figure S13:**
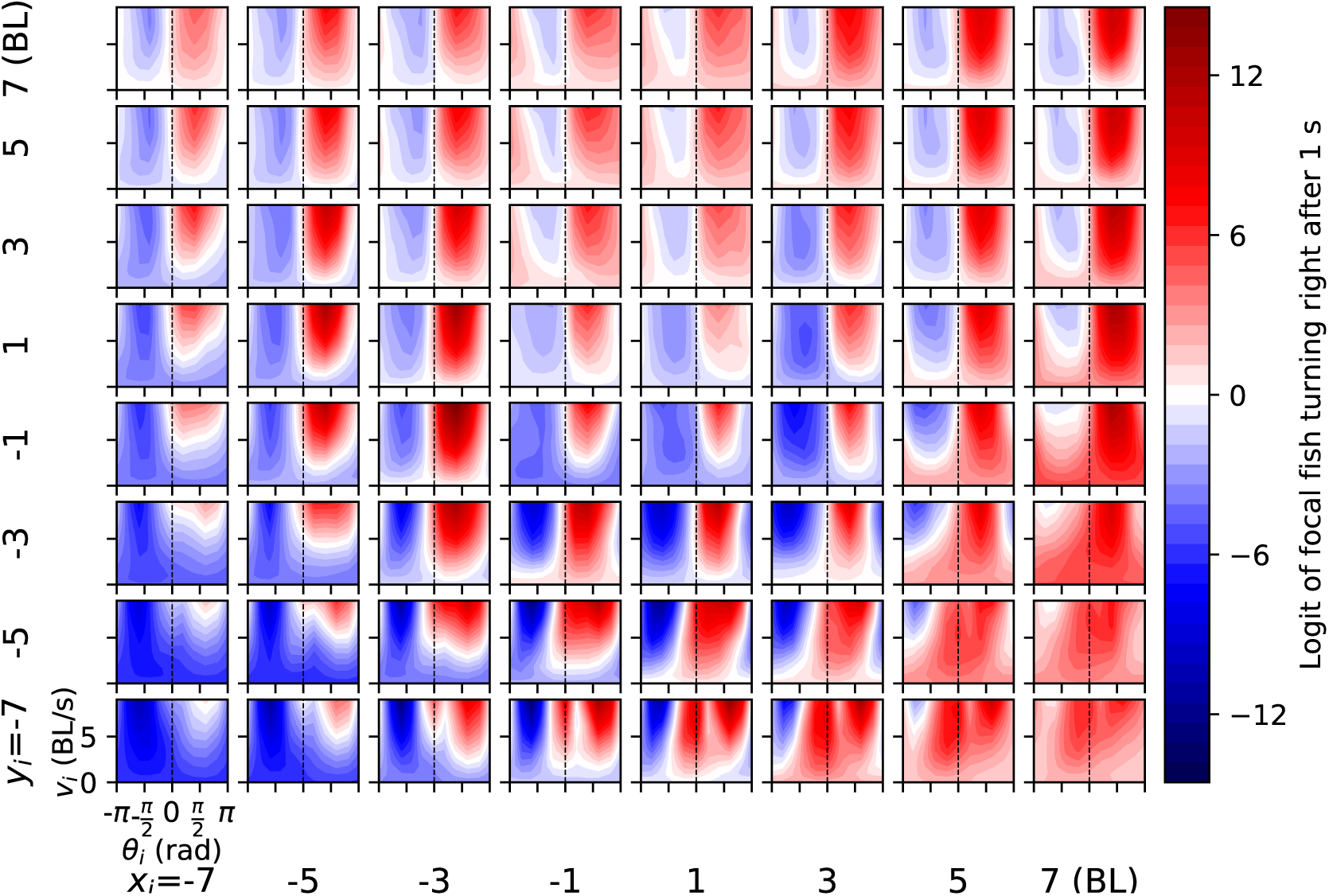
Properties of interaction between a pair of fish in the collective when the focal fish is in the midst of a right turn, focal normal acceleration fixed to *a*_⊥_ = 100 BL/s^2^.

**Figure S14:**
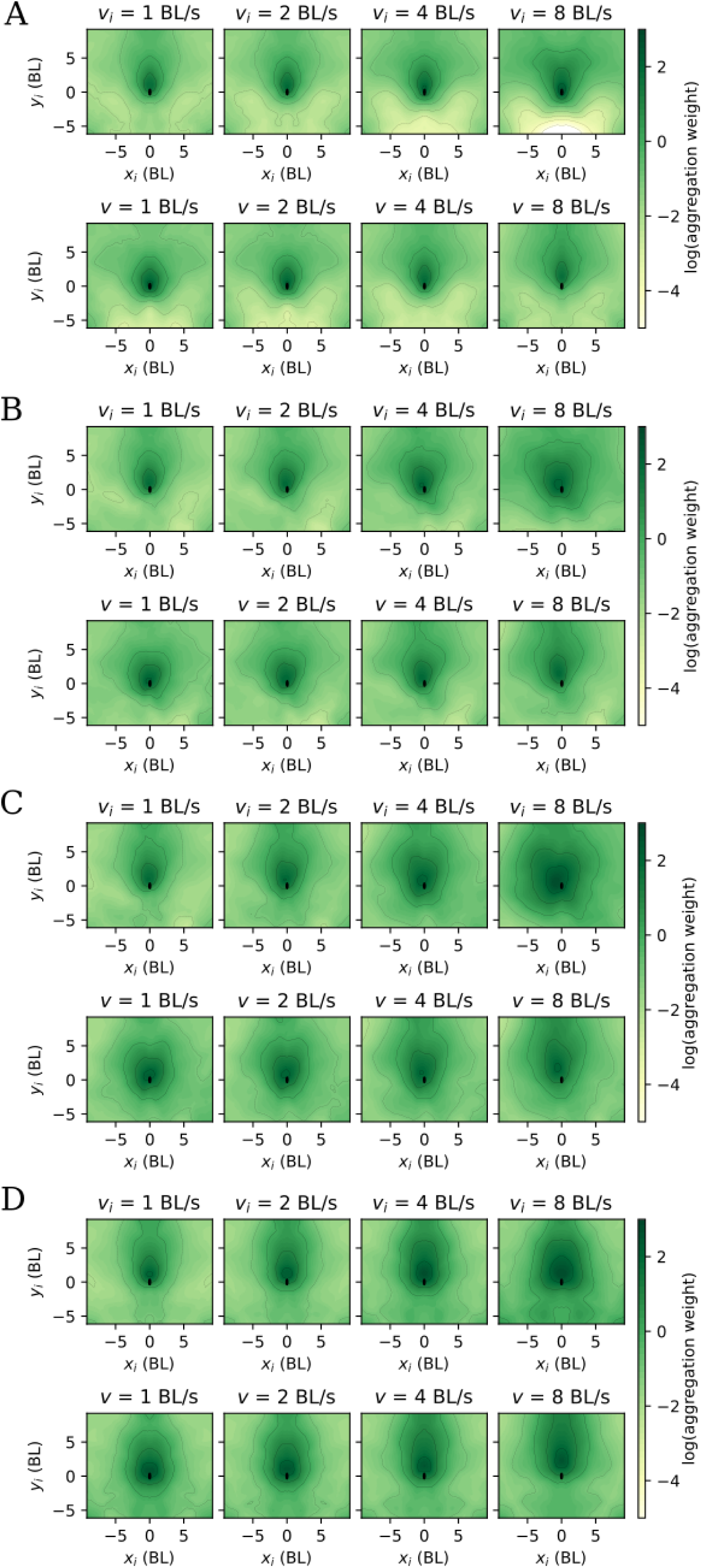
Aggregation with information about relative orientation. Same as Figure 4, but when the attention subnetwork is trained with the relative orientation of the neighbour, in addition to the variables used in the main text. **A** The neighbour is parallel (at 0 degrees) to the focal. **B** The neighbour is at 45 degrees (towards the right) with the focal, **C** the neighbour is perpendicular (90 degrees) and pointing to the right of the focal. **D** The neighbour is antiparallel (180 degrees) to the focal.

